# Aurora A mediated new phosphorylation of RAD51 is observed in Nuclear Speckles

**DOI:** 10.1101/2023.08.11.552966

**Authors:** Mohamad Alaouid, Parfait Kenfack Ymbe, Vanessa Philippot-Ménil, Gwennina Cueff, Alexandre Demeyer, Damien Marquis, Nizar Ayadi, Fabrice Fleury, Houda Benhelli-Mokrani

## Abstract

To maintain its genome integrity, the cell uses complementary and orchestrated processes, among which is the DNA Damage Response. Pre mRNA maturation is an essential step of the DNA damage response that provides an adapted proteome in order to face genotoxic stress. We describe here a new phosphorylation of the RAD51 recombinase, on its Ser97 residue. This new Aurora A mediated RAD51 phosphorylation modulates its *in vitro* activity evaluated by D-loop and polymerization assays. Using recombinant proteins, we show that RAD51 is an RNA binding protein and that the Ser97 phosphorylation modulates its RNA binding affinity *in vitro*. Using a specifically generated antibody we revealed that this Ser97 phosphorylation is correlated with RAD51 localization into the RNA maturation membrane less organelles, Nuclear Speckles. We describe here for the first time the presence of RAD51 DNA repair factor within the Nuclear Speckles, raising the hypothesis of a possible role for RAD51 in splicing modulation. This point is of particular interest in the context of splicing profiles modulations associated with radio and/or chemoresistance.

## INTRODUCTION

Repair of DNA damages is essential for genomic stability and a miss-regulation of this essential DNA metabolism can cause cancer. The cell response to DNA damages includes a cell cycle arrest in G1/S or G2/M, depending on the cell cycling and the DNA damage, to provide the time needed to repairing its genome before cycling again ^1^. In the case of a large amounts of damages, the cell undergoes apoptosis and this is the principle used in chemo- and radiotherapies to kill cancer cells (for review ^2^). Aurora A is a well-known kinase implicated in mitosis orchestration and its kinase activity has been described as diminishing after DNA damages ^3^.

The RAD51 recombinase is implicated in the strand exchange mechanism during the DSB repair by the Homologous Recombination (HR) pathway. In the absence of DNA Damage (DD), RAD51 is predominantly cytoplasmic and translocates to the nucleus during the DNA Damage Response (DDR) to manage HR repair ^4 5^. As it needs the undamaged sister chromatid as a template, the HR repair pathway occurs mainly in the late S, G2 phases of the cell cycle ^6 7^. However, it has been documented that HR repair can also occur during G1 and early S phases, and in this case, the undamaged template used for the repair could be the homologous chromosome or an RNA transcript ^8^. These mechanisms of sister chromatid-less HR repair may induce a Loss Of Heterozigosity (LOH) which is frequently observed in cancer development ^9 10^. Thus, RAD51 localization and activity must be controlled and finely regulated in order to maintain genomic stability. For that purpose, RAD51 is post-translationally modified by multiple phosphorylations which affect its auto-polymerization, DNA binding capacities, and D-loop formation ^11–13^.

When considering the DDR, we usually direct our thoughts to the cascade of post-translational modifications (PTM), and in particular the phosphorylation which is the most studied. These PTM thus regulate protein/protein and protein/DNA interactions in the context of chromatin remodeling and DNA repair. This view is completed by the consideration of protein variant adaptation. Indeed, the response to DD is also regulated at the post-transcriptional level by a modulation of pre-mRNA maturation making possible the production of proteins isoforms ^14 15^. Active splicing machinery is essential during the DDR, allowing the cell to repair its DNA damages and maintain its genomic stability. That is why alternative splicing deregulation could be implicated in carcinogenesis as supposed by many studies showing abnormal alternative splicing profiles in cancer cells compared to the non-pathological cells ^16,17^. We know from Pederiva’s work that inhibiting splicing before DNA damage induction impedes the recruitment of DNA repair factors to the damaged sites ^18^. We must consider that the splicing modulation occurring after DNA damages takes part of the DDR itself. In this sense, some well-known players in the signaling cascade that allow DNA damage repair have been described as also being implicated in splicing modulation in the context of the DDR. Thanks to the work of Katzenberger, we know that ATM and ATR kinases and their respective targets, chk2 and chk1 checkpoint kinases, are essential for the splicing modulation of the TAF1 factor after DNA damage ^19^. The work of Matsuoka, consisting of identifying ATM and ATR targets after IR, using the SILAC approach, revealed news pathways activated during the DDR. Of them, there is the RNA metabolism where many proteins implicated in pre mRNA maturation are phosphorylated by ATM or ATR ^20^. Even the number of studies showing a link between DNA repair and RNA splicing is growing, (for review ^21^), we are still in the identification process for the connections between these two branches of the cell response to DNA damage. Here, we identified an Aurora A mediated new phosphorylation of RAD51 that is located into nuclear structures that are not γ−H2AX DNA repair foci. We showed that this new phosphorylation of RAD51 is located into Nuclear Speckles, the RNA maturation sites. In this work, we also show that RAD51 is an RNA binding protein and that this Ser97 phosphorylation affects its *in vitro* RNA binding affinity.

## MATERIAL AND METHODS

### RAD51 recombinant protein production

WT and RAD51 mutants (S97D phosphomimetic), (S97A non phosphorylable) were produced according to the previously described protocol ^12^. Briefly, pet15b plasmids encoding His-tagged RAD51, WT and (S97A, S97D) mutated versions were used in the E coli BL21 DE3 strain at 37°C. After IPTG induction, the bacteria were lysed with Tris HCl 50 mM, NaCl 500 mM, glycerol 10%, β-mercaptoethanol 5 mM, imidazole 5 mM and recombinant RAD51-proteins were purified on a NiNTA resin (#P6611, Sigma, St Louis, MO, USA) using the washing buffer: Tris HCl 50 mM, NaCl 500 mM, glycerol 10%, β-mercaptoethanol 5 mM, (and increasing imidazole concentrations).

A dialysis step was then added (dialysis buffer: Tris HCl 50 mM, NaCl 500 mM, glycerol 10%, EDTA 1 mM, DTT 1 mM). A Bradford assay was used to determine protein concentrations and SDS-PAGE followed by Coomassie staining were used to check the purity of the protein fractions. *in vitro* phosphorylation/phosphorylated site MS identification.

### *In vitro* phosphorylation assay and MS analysis

Recombinant Aurora A active protein was purchased at SIGMA (A1983). 5 µg of recombinant RAD51 were phosphorylated *in vitro* at 37°C for 30 minutes in 50 mM Tris HCL pH=7,5, 10mM MgCl2, 50uM DTT, 50uM ATP, and 200 ng of Aurora A enzyme. Phosphorylation products were separated using 12% acrylamide gel electrophoresis.

Proteins were colored with the BIO-SAFE Coomassie stain reagent (BIORAD, 1610786) according to the manufacturer’s recommendations. Gels slices were maintained in 1% acetic acid solution at room temperature before their digestion and analysis. MS/ORBITRAP analysis was performed by the 3P5 core facility (Institut Cochin, Paris).

### D-loop assay

As described previously in Chabot et al ^12^: 100-ssDNA labeled IRD700 (1 μM) (Integrated DNA Technologies, Coralville, IA, USA) was incubated with 0.5 μM RAD51 in 10 μL of standard reaction buffer containing 20 mM Tris HCl pH=8, 1 mM ATP, 1 mM DTT and 1 mM CaCl2 at 37 °C for 20 min. Supercoiled complementary plasmid pPB4.3 DNA was added at a final concentration of 200 μM (in bp), to initiate the strand exchange reaction. After an incubation time of 30 min at 37 °C, the reaction was stopped by the adding 10 mM of Tris HCl pH=8, 10 mM MgCl2 and 1% SDS (final concentrations) to the reaction. 1 mg/mL of proteinase K was used to deproteinize the reaction for 15 min at 37 °C. The reaction products were then separated by electrophoresis on a 1% agarose gel, in 0.5× TAE buffer (20 mM Tris, 10 mM acetic acid and 1 mM EDTA) at 100 V for 2 h. Detecting the IRD-700 dye with the 700 nm infrared fluorescent detection channel of the Odyssey Infrared Imager (LI-COR Biosciences) allowed us to detect and quantify the ssDNA and D-loop structures.

### DNA /RNA binding activity

Blitz technology was used with 100 mM of 5’ biotinylated 33 nt DNA or RNA bound on a streptavidin coated biosensor. DNA and RNA sequences are given below. WT and mutants RAD51 recombinant proteins were diluted at the indicated concentration in PBS buffer 1× and ATP 2 mM. Three steps kinetics were made. The buffer baseline was measured for 10 sec, then the association step between ssDNA or ssRNA and RAD51 at different indicated concentrations from 0,5 to 4 µM was monitored for 200 sec. Finally, the dissociation step of the bound RAD51 was monitored for 200 sec in buffer. Tip was regenerated in NaOH (50 mM) twice for 40 sec before reuse. Nucleic acids sequences:

33m DNA: 5’ TCCTTTTGATAAGAGGTCATTTTTGCGGATGGC 3’

33m RNA: 5’ GCCAUCCGCAAAAAUGACCUCUUAUCAAAAGGA 3’

### Antibodies

Rabbit antiPSer97 RAD51antibody was specifically produced by Genecust. Mouse anti-RAD51 antibody is from Invitrogen, ref: MA5-14419. Rabbit anti-RAD51 antibody is from Invitrogen, ref: MA5-32789. Anti-tubulin antibody is from Invitrogen, ref: MA1-19162. Anti-histone H3 antibody is from Invitrogen, ref: PA5-16183. Anti-gH2AX antibody is from Merck Millipore, ref: JBW301. Goat anti rabbit secondary antibody is from LI-Cor, ref: D00804-07, Goat anti mouse secondary antibody is from LI-Cor, ref: D00804-13.

### Immunofluorescence labelling/Microscope acquisitions

Cells were seeded on a 1cm diameter glass. At the indicated time, the cells were washed twice with PBS then fixed for 15min in 3.7% formaldehyde in PBS. After 3 PBS washes, slides were kept at 4°C before immunolabelling. Briefly, cells were permeabilized in triton 0.01%, washed with PBS 1% BSA and incubated with primary antibodies diluted in PBS 1% BSA for 2 hours. After 3 washes, the cells were incubated with an Alexa conjugated secondary antibodies for 30 min. After 3 washes, the cells were mounted in Prolong DIAMOND+DAPI mounting medium (Thermofisher, P36962). Image acquisitions were made on a NIKON epifluorescence microscope and on a NIKON confocal microscope. Confocal microscopy acquisitions were made at the IBISA MicroPICell facility (Biogenouest), member of the national France-Bioimaging infrastructure supported by the French national research agency (ANR-10-INBS-04). Image analysis and figures realization were made using Fiji and the Figure J plugin. Colocalization studies were performed in accordance with the recommendations of Dunn Practical guide to evaluation of colocalization in biological microscopy ^22^. JACoP tool on Fiji was used to calculate the Pearson’s Correlation Coefficient (PCC). This analysis was performed in a cell-by-cell ROI (regions of interest) PCC determination. Two signals are colocalized when a PCC value is positive, and they are anti-colocalized which means an exclusion when the PCC value is negative. A null PCC value means a random distribution of the two signals. We used the Zinchuk et al., 2013 methodology to interpret PCC colocalization values. It proposes a classification of 5 degrees of colocalization: PCC from 0 to 0.12: very weak/ PCC from 0.13 to 0.39: weak, PCC from 0.4 to 0.6: moderate, PCC from 0.49 to 0.84: strong, PCC from 0.85 to 1: very strong ^23^. As recommended by the Mcdonald et al. work, the comparative analysis of two conditions, (Ctl and plaB treatment for example) was made using a unilateral t-test ^24^.

### Cell culture and treatments

The human basal-like TNBC cell line, HCC1806, and HeLa were grown in 100 mm x 20 mm dishes (for sub-fractionation experiments) or six-well plates (for total extract experiments) at 37 °C in a humidified atmosphere containing 5% CO2. RPMI 1640 medium 1X (Life Technologies, Carlsbad, CA, USA) and DMEM medium 1X (Dulbecco’s Modified Eagle Medium) were used as growth media for HCC1806 and HeLa, respectively. Each growth medium was supplemented with 10% fetal bovine serum and 1% penicillin /streptomycin (REF15140-122, Gibco). Cells were treated with 10 µM camptothecin (C9911-100MG, Sigma), 5 µM Pladienolide B (SC-391691, Santa Cruz Biotechnology) for 4 hours, or 50 nM alisertib (cat.# HY-10971/CS-0106, Medchemexpress®) for 24h to inhibit Aurora-A activity (alisertib was dissolved in DMSO and then stored at −20°C). The cells then underwent protein total extract or sub-fractionation experiments as described below.

### Total cell extracts

HeLa and HCC1806 cells were cultured in six-well plates. After different treatments, the cells were washed twice with room temperature PBS (Gibco, REF 14190-094). Proteins were extracted by treating the plates with RIPA buffer (25 mM Tris HCl pH=7.6, 150 mM NaCl, 1% NP-40, 1% sodium deoxycholate, 0.1% SDS) (#89900, Thermo scientific), All extraction steps were carried out on ice.

### Subcellular fractionation

Following application of the desired treatments, the cells were rinsed twice with room temperature PBS, trypsinized, and collected in 1 ml Eppendorf tubes. They were washed again with buffer A (10 mM HEPES pH=7.9, 1.5 mM MgCl2, 10 mM KCl, 0.5 mM DTT), and centrifuged for 5 minutes at 1000 rpm. They were then homogenized in buffer A supplemented with (0.1% NP-40, 1mM phosphatase inhibitor cocktail3, 1mM PMSF, 1/100 protease inhibitor cocktail) and incubated on ice for 10 minutes. The cytoplasmic fractions correspond to the supernatants obtained after 5 minutes of centrifugation at 5000 rpm. They were then washed twice with 1 ml of buffer A and centrifuged at 5000 rpm for 5 minutes. Following, they were agitated for 15 minutes in buffer B (20 mM HEPES pH=7.9, 25% glycerol, 0.42 M NaCl, 1.5 mM MgCl2, 0.2 mM EDTA, 0.5 mM DTT) complemented with (1 mM phosphatase inhibitor cocktail3, 1 mM PMSF, 1/100 Protease Inhibitor Cocktail) and centrifuged at 14000 rpm for 15 minutes. The supernatants collected correspond to the nuclear fractions. The pellets were then washed twice with 1 ml of buffer B and centrifuged at 14,000 rpm for 15 minutes. Depending on the pellet volume, 50 to 75 μl of buffer B with (1 mM phosphatase inhibitor cocktail3, 1 mM PMSF, 1/100 Protease Inhibitor Cocktail) were added. Finally, the pellets were sonicated at 70% amplitude for 1 min, with a sonication pulse rate of 1 second on/ 1 second off. The fractions obtained represent the proteins bound to chromatin. Throughout the process, the samples were stored on ice and the centrifugations were conducted at 4 °C.

### Cell transfections

HeLa and HCC1806 cells were transfected for the time indicated (24 or 48h) with purified Aurora-A plasmid, or indicated siRNA, using Lipofectamine 2000 (Invitrogen) according to the manufacturer’s instructions with a ratio of DNA (or RNA)/lipo2000 of 1/1. Scramble and RAD51 siRNA were purchased from Origene technologies, ref: SR321568.

### Western blot assay

Proteins from cell lysates were separated on 12% SDS-polyacrylamide gels by electrophoresis. For western blot analysis, we transferred the proteins onto nitrocellulose membranes (GE Healthcare, Little Chalfont, UK) using the wet transfer method for 1 hour at 250 mA (4 °C). Membranes were blocked for 30 min at room temperature with agitation using 5% non-fat dry milk (w/v) in PBST (1x PBS, 0.1% Tween 20) (v/v). They were rinsed twice for 5 min in PBST, after which the membranes were incubated with the primary antibody overnight with agitation at 4 °C. Then, they were incubated with the secondary antibody for 30 min in the dark at room temperature and washed (3X, 5 min) with PBST. The Odyssey infrared imaging system (LI-COR Biosciences, Lincoln, NE, USA) was used to visualize immunoreactive bands. Image Studio Lite software version 5.2 (LI-COR) was used to quantify and normalize protein of interest signals to loading controls (α-tubulin and H3).

### Dot blot assay

Recombinant RAD51 proteins were spotted on nitrocellulose membrane (GE Healthcare, Little Chalfont, UK). Membranes saturation, antibodies incubation and scanning were performed in the same way as for western blot assays.

### Statistical analysis

A paired Student’s t-test was used in the statistical analysis using Microsoft Excel 2016. Statistical significance was considered to be present when p ≤ 0.05 (*), p ≤ 0.01(**), p ≤ 0.001(***), or not significant (ns) when p > 0.05. Each error bar represents the standard deviation.

### Cell synchronization

HeLa cells were grown in a T75 flask until they were around 35–40% confluent. After that, 2 mM of thymidine (Sigma, T9250-1G) was added, and the cells were incubated for 18 hours. Following this, thymidine was removed, and the cells were washed twice in fresh DMEM. After that, fresh DMEM supplemented with 24 µM deoxycytidine was added for 11 hours. The second blockade was obtained by incubating cells in thymidine 2 mM for 14 h. Then the thymidine was removed, and the cells were washed twice in fresh DMEM. The second turn of 24 µM of deoxycytidine in DMEM was then done, followed by cell harvesting after 0h −2h −6h −8h −10h and12h.

After ethanol fixation and IP staining, cells were passed in a FACS. Cell cycle analysis were done using FlowJo ® software.

### RAD51 immunoprecipitation

HCC1806 cells were collected and lysed with Pierce® IP lysis buffer (87787, Thermo), which contains: (25 mM Tris -HCl pH 7.4, 150 mM NaCl, 1% NP-40, 1 mM EDTA, 5% glycerol) and supplemented with 1 mM phosphatase inhibitor cocktail3 (Sigma, P0044), 1 mM PMSF, 1/100 protease inhibitor cocktail (Sigma, P8340). The lysate was then clarified by centrifugation (14,000 rpm, 15 min), and divided into two tubes, one of which received 100 µg/ml of RNase A (EN0531, Thermo Fisher), and the other of which served as a control, the tubes were then incubated at 37 °C for 20 minutes. After that, one milligram of proteins was then incubated overnight with 10 µg of anti-pSer97 RAD51 antibody (Gene Cast). The resulting immune complex was then incubated for 1 h with 40 µl of protein A/G plus agarose beads (sc-2003, Santa Cruz). After this, the beads were washed four times with IP lysis buffer, and the proteins were then eluted by heating the beads for 5 min at 95 °C in 50 μl of 2x SDS buffer (Laemmli 2x ref S3401-1VL, Sigma), Finally, IP fractions were subjected to electrophoresis.

### Computational mutagenesis

The 3D structure of RAD51 used is a nucleofilament consisting of a trimer of HsRAD51 and a dsDNA; it is available in the Protein Data Bank under accession code 5H1C.

The analysis and the 3D rendering are achieved on the open-source Pymol 2.5 software (https://github.com/schrodinger/pymol-open-source).

The mutagenesis of HsRAD51 from S97 to S97A or S97D was performed using the !Mutagenesis” tool in Pymol.

### Polymerization assay

20 μM HsRAD51 was incubated in 20 mM HEPES pH=7.5, 150 mM KCl, 20 mM DTT for 30 min at 4°C. HsRAD51 complexes were cross-linked by incubation with 0.5 or 10 mM of BS3 (Sigma-Aldrich) at 22°C for 30 min. BS3 allows reticulation between different amines located on the residues side chains and terminal amins functions of RAD51 monomeres. Cross-linking was terminated by adding 80 mM Tris pH=8.0 for 15 min at 22°C and analyzed with SDS-PAGE. The gels were colored with Coomassie-Blue staining and the Odyssey infrared imaging system (LI-COR Biosciences, Lincoln, NE, USA) was used to visualize immunoreactive bands. Image Studio Lite software version 5.2 (LI-COR) was used to quantify and analyze band intensity signals.

## RESULTS

### Aurora A kinase phosphorylates RAD51 *in vitro* within its subunit rotation motif

Recombinant HsRAD51 protein was used to perform an Aurora A kinase mediated *in vitro* phosphorylation. Figure 1A shows a Coomassie-Blue stained SDS-PAGE, with a shift in the last well corresponding to the RAD51 with ATP and Aurora A kinase condition. Differential MS analysis in comparison with the unphosphorylated form allowed us to identify a unique phosphorylated site on the Serine 97 (see Supplemental Figure 1). This residue is located in the linker region of RAD51, between the N-terminal domain (Nter) and the Core domain (Core), within the Subunit Rotation Motif (SRM) as shown in figure 1A. In this same figure, the sequence alignment of RAD51 linker regions from different species shows the conservation of the phosphorylated residue, represented in bold. The 3D structure of HsRAD51 was used to visualize the Ser97 residue within the RAD51 nucleofilament. We can see in figure 1B that this residue remains exposed at the surface of the nucleofilament and thus, is available for potential modulation of the interactions of RAD51 with its partners. Recombinant S97A and S97D, respectively non phosphorylable and phosphomimetic mutants, were produced and used to test the effect of this phosphorylation on RAD51 activity. Figures 1B2 and 1B3 show the S97A and S97D mutations respectively. The side chains of these mutants are free like that of the Ser of the WT RAD51. We used these recombinant mutants RAD51 to explore their molecular phenotype and predict the impact of the phosphorylation of the Ser97 residue on RAD51 activity.

**Figure 1.**
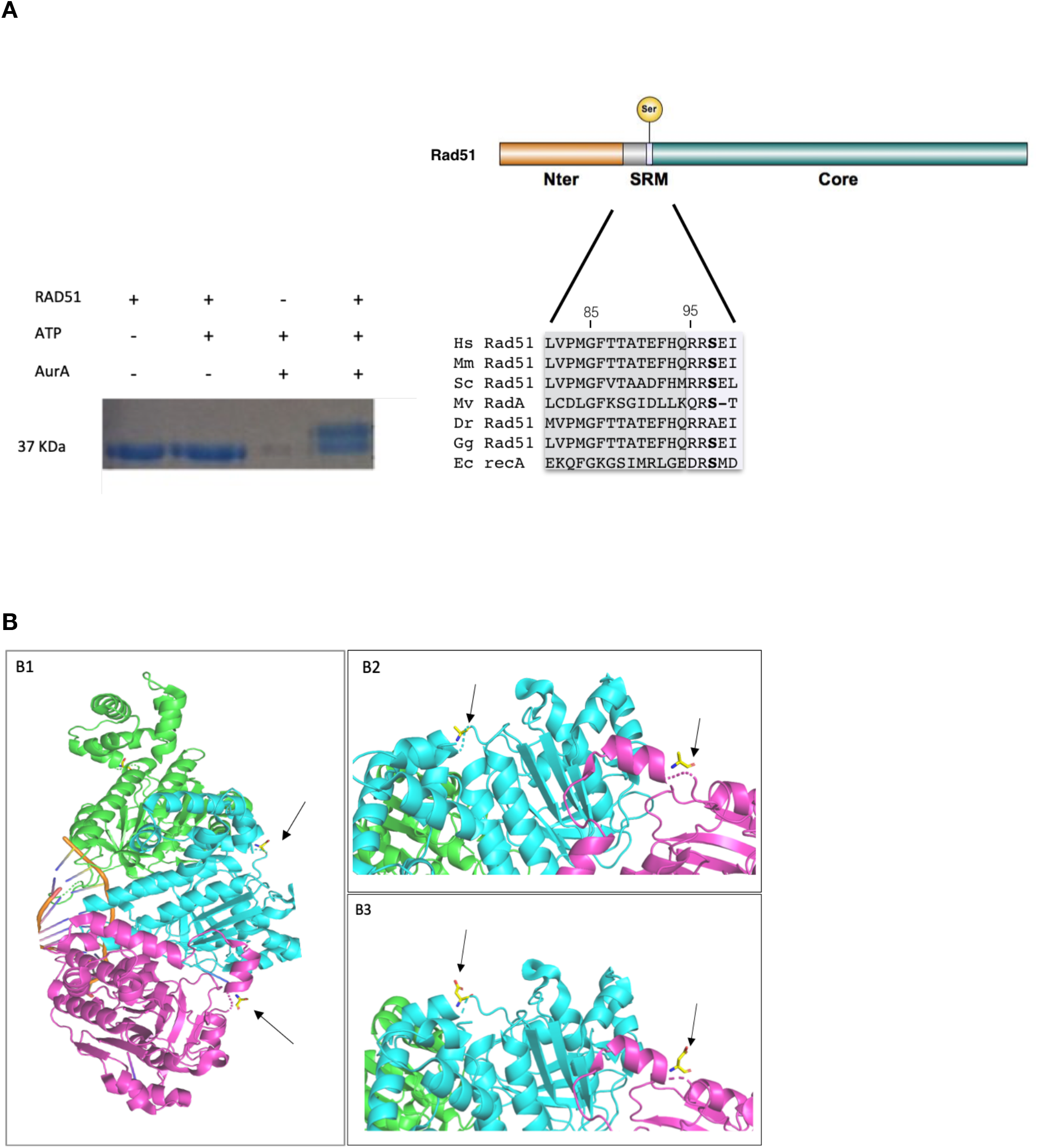
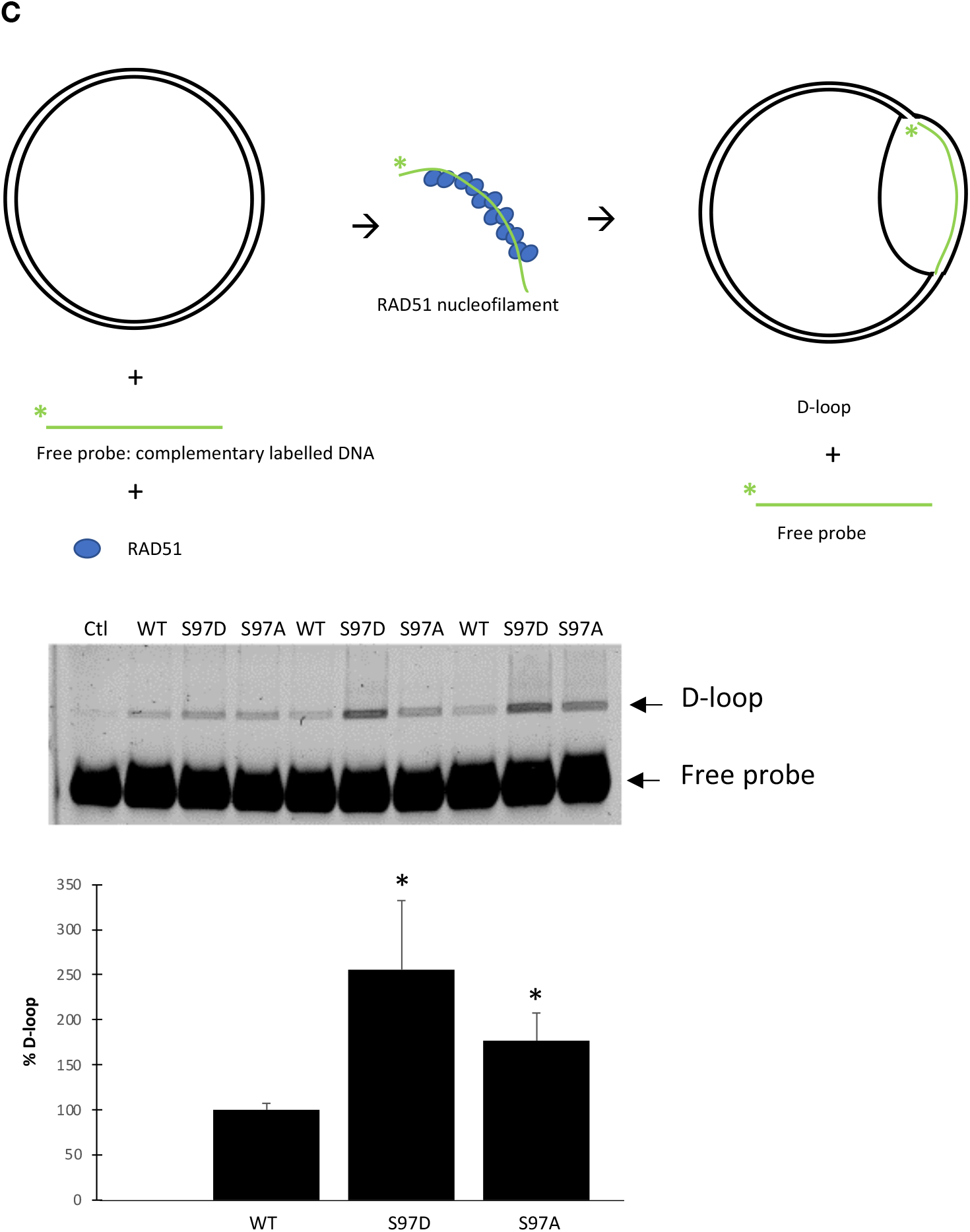
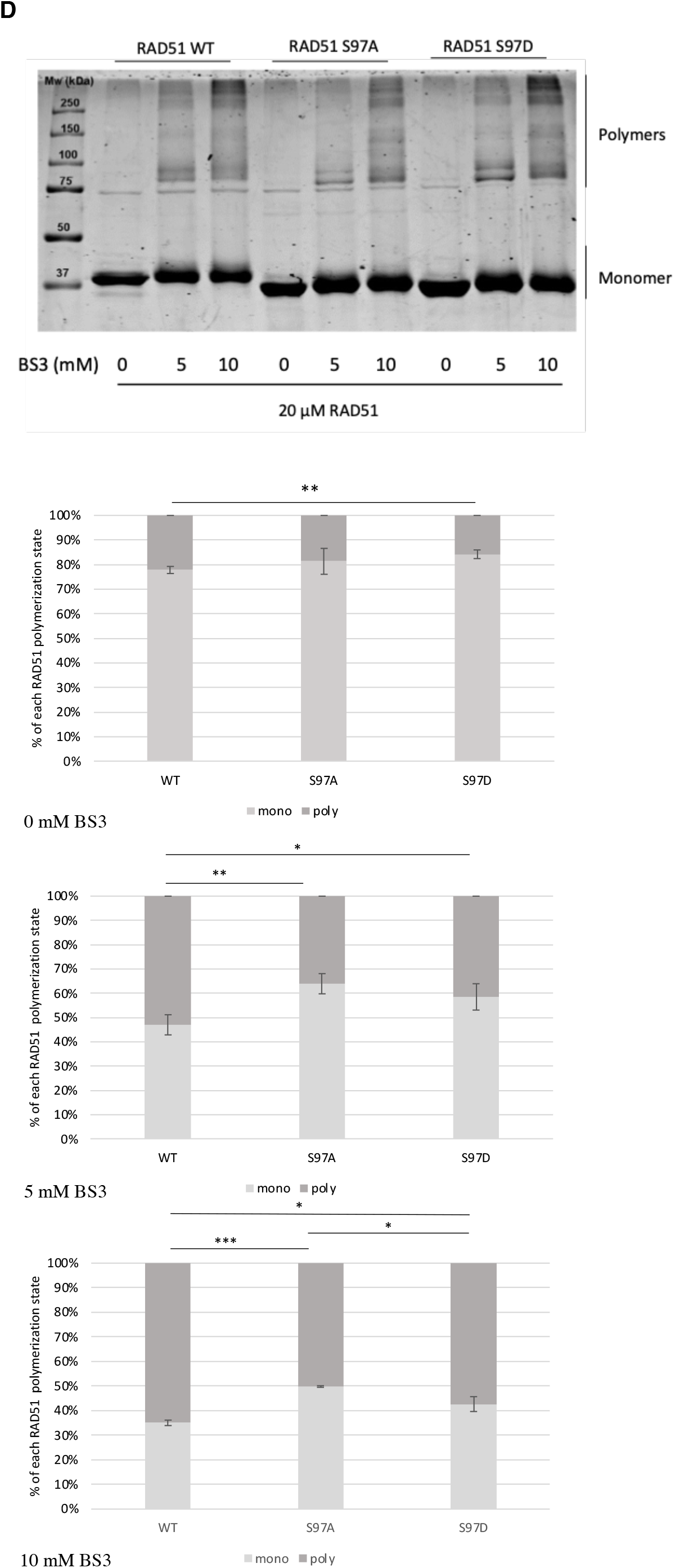
**A: Aurora A kinase phosphorylates RAD51 *in vitro* on its Ser97 residue** Coomassie blue stained SDS-PAGE after *in vitro* phosphorylation of recombinant RAD51 by Aurora A kinase. Within the Subunit Rotation Motif, the Ser97-RAD51 residue is well conserved among species. The AurA-phosphorylated residue is located within the Subunit Rotation Motif (SRM) of RAD51 Nter: N terminal domain, Core: Core domain Sequence alignment of RAD51 Linker Region (81 to 99). Light gray background corresponds to the Subunit Rotation Motif (95 to 99). S in bold corresponds to the residue phosphorylated in HsRAD51. Hs: Homo sapiens, Mm: Mus musculus, Sc: Saccharomyces cerevisiae, Mv: Moloney virus, Dr: Danio rerio, Gg: Gallus gallus, Ec Escherichia coli. **B: Prediction of the effect of Ser97-phosphorylation on RAD51 structure** The 3D structure of HsRAD51 WT allows us to observe Ser97 which is located in a random coil on the linker region of each monomer precisely in the SRM (B1). By zooming in on a part of the nucleofilament where Ser97 has been mutated to Ala, S97A mutation (B2) or to Asp, S97D mutation (B3), we can see that the side chains of these residues are free like that of Ser97. Arrows indicate the residues located on the 97 position (Ser, Ala or Asp in the B1, B2 and B3 pictures respectively). **C: RAD51 phosphorylation mimetic on Ser97 affects its D-loop activity** The D-loop test was used to evaluate the D-loop formation capacity of the of WT, S97A and S97D-RAD51 recombinant proteins. Top figure: a scheme of the experimental D-loop assay. The intensities of the free probe and the D-loop were quantified by an IRD700 scan, then used to evaluate the D-loop formation. The activity of the WT-RAD51 was set as 100% and used for S97A and S97D activity standardization. T-test with alpha=0,05 was used: p-value WT *vs* S97D=0,025, p-value WT *vs* S97A=0,014 (n=3 experiments). **D: RAD51 phosphorylation mimetic on Ser97 affects its polymerization rate** WT, S97D and S97A RAD51 recombinant proteins were used to evaluate the effect of RAD51 phosphorylation on its polymerization, by the *in vitro* polymerization assay. RAD51 was incubated with or without BS3 reticulating agent, then used to perform an acrylamide gel electrophoresis and a Coomassie blue staining. Intensities of n=3 experiments were quantified using a Lycor scan and used for statistical analysis (t-test with alpha= 0,05 was used: without BS3, Wt *vs*S97D, p-value= 0.008. For the 5mMBS3 condition, p-value WT *vs* S97A=0.007, p-value WT *vs* S97D= 0.04. for the10mM BS3 condition, p-value WT *vs* S97A=2.3 10 ^−5^, p-value WT *vs* S97D= 0.015, p-value S97D *vs* S97A= 0.016).

### The S97D RAD51 phosphomimetic mutant shows increased *in vitro* strand invasion capacity

To evaluate the effect of the Ser97 phosphorylation on RAD51 activity, we performed a D-loop assay experiments and evaluated the strand invasion capacity of WT, S97A and S97D-RAD51 recombinant proteins. This assay allows to quantify the RAD51 enzymatic activity which favors the strand invasion of a free labelled probe within a plasmid containing the same sequence, to form a displacement loop, (refer to figure 1C illustration of this technique). We observed that the phosphomimetic mutant, S97D has an increased D-loop formation ability. Relatively to the D-loop activity of the WT RAD51, set as 100%, the S97D-RAD51 activity was more than two-times higher. This difference was statistically significant. We noticed that the S97A-RAD51 mutant also had increased activity, about 50% more than the WT one. These results highlight the importance of the Serine 97 on RAD51 activity.

### The S97D RAD51 phosphomimetic mutant has a decreased polymerization rate

In order to evaluate the impact of the Ser97 mutations on RAD51 protein/protein interactions, we performed a polymerization assay without the presence of DNA. RAD51 was incubated at different concentrations with or without the amine reticulating agent BS3, then loaded in an acrylamide gel. (BS3 is used here to stabilize, proteinic interactions. Refer to the materiel section for more details). The gels were stained with Coomassie-Blue and the band intensities of monomeric or polymeric RAD51 were quantified and analyzed. We can see, in Figure 1D that the Ser97 mutations affects the RAD51 polymerization state. Both S97A and S97D mutants showed a statistically significant decreased ability to polymerize, meaning that RAD51/RAD51 interaction without nucleic acids are affected by the Ser97 Phosphorylation. These results highlight the importance of the Ser97 residue in RAD51 self-association ability, in accordance with the already described role of the SRM (that includes the Ser97 residue) on RAD51 polymerization.

### RAD51 Ser97 is phosphorylated *in cellulo*

We generated a specific antibody directed against the phosphorylated Ser97 form of RAD51 (PSer97-RAD51). This rabbit raised phosphospecific antibody was validated by different complemental techniques presented in the figure 2A panel. Using dot-blot assay, the phosphospecific antibody was tested for its ability to recognize two peptides corresponding to the RAD51 SRM region, with only one of them phosphorylated on its Ser97. The results are presented in the figure 1A.1 and show that the generated antibody recognizes only the peptide with the phosphorylated Ser97. Peptide competition assays were also used in order to validate the phospho-antibody specificity. Results are shown in figure 2A.2. The PSer-RAD51 signal almost disappeared after PSer-RAD51 peptide competition assay, while it is maintained after the competition with the Ser-RAD51 non-phosphorylated peptide. Bovine Intestine Phosphatase was also used to treat half of a membrane, before its immunoblotting was performed. We observe in the figure 2A.3, that on the BIP treated part of the membrane, we see only a little remaining PSer-RAD51 signal and more non-specific signals. This experiment prove that the signal detected by this antibody is a phosphorylation. BIP was also used directly after Aurora A mediated *in vitro* phosphorylation of recombinant RAD51 protein, before performing a western blot. Results are shown in the figure 2A.4, in which the comparison of the wells n°5 and 10 allows us to see that the BIP treatment strongly diminished the signal detected by the anti-PSer97-RAD51 antibody. We also used siRNA mediated RAD51 depletion to test the antibody specificity. The results from n=3 experiments, illustrated in figure 2A.5 showed that RAD51 siRNA induced a 63% diminution of RAD51 protein level and a 51% diminution of PSer97-RAD51 level.

**Figure 2.**
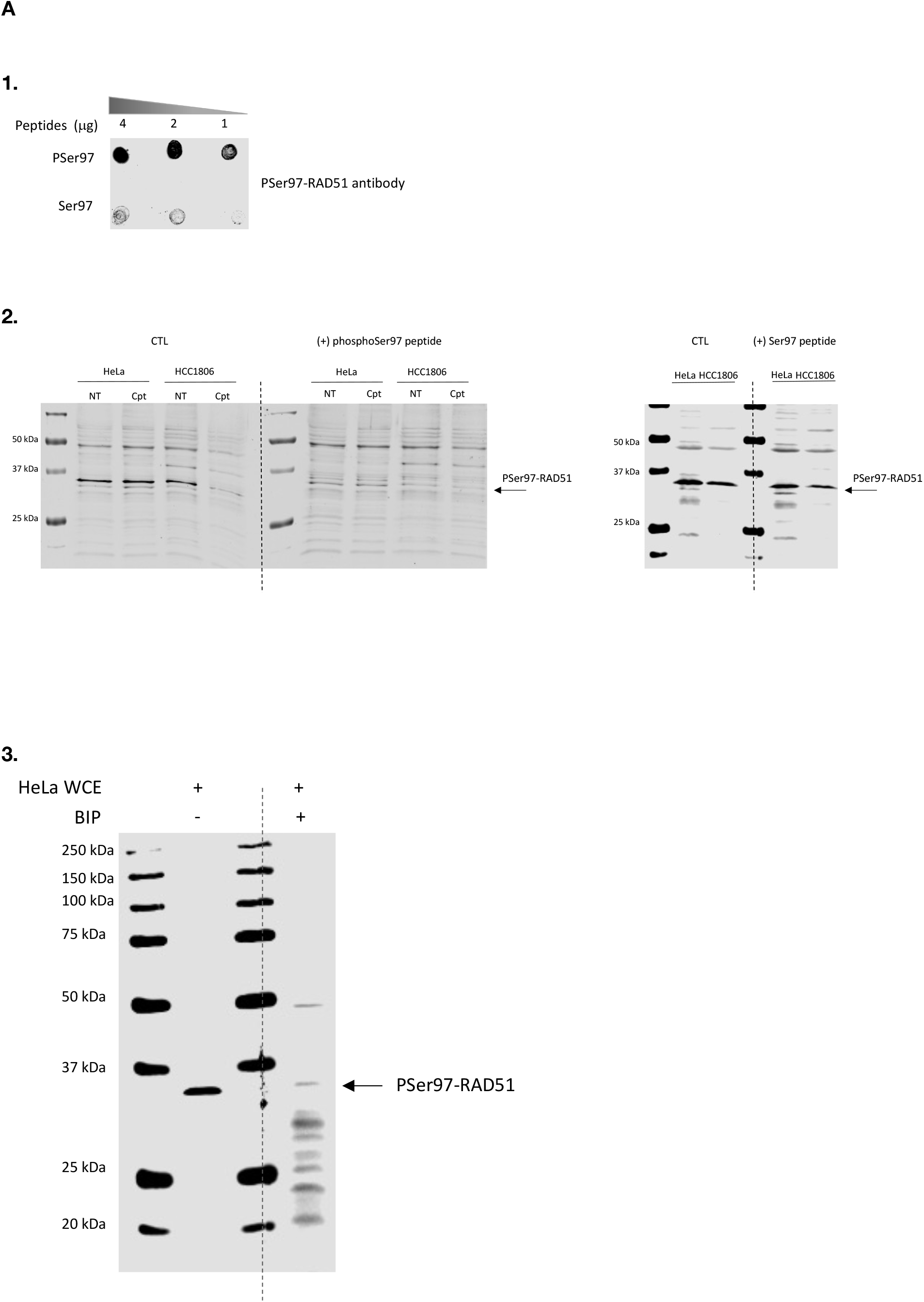
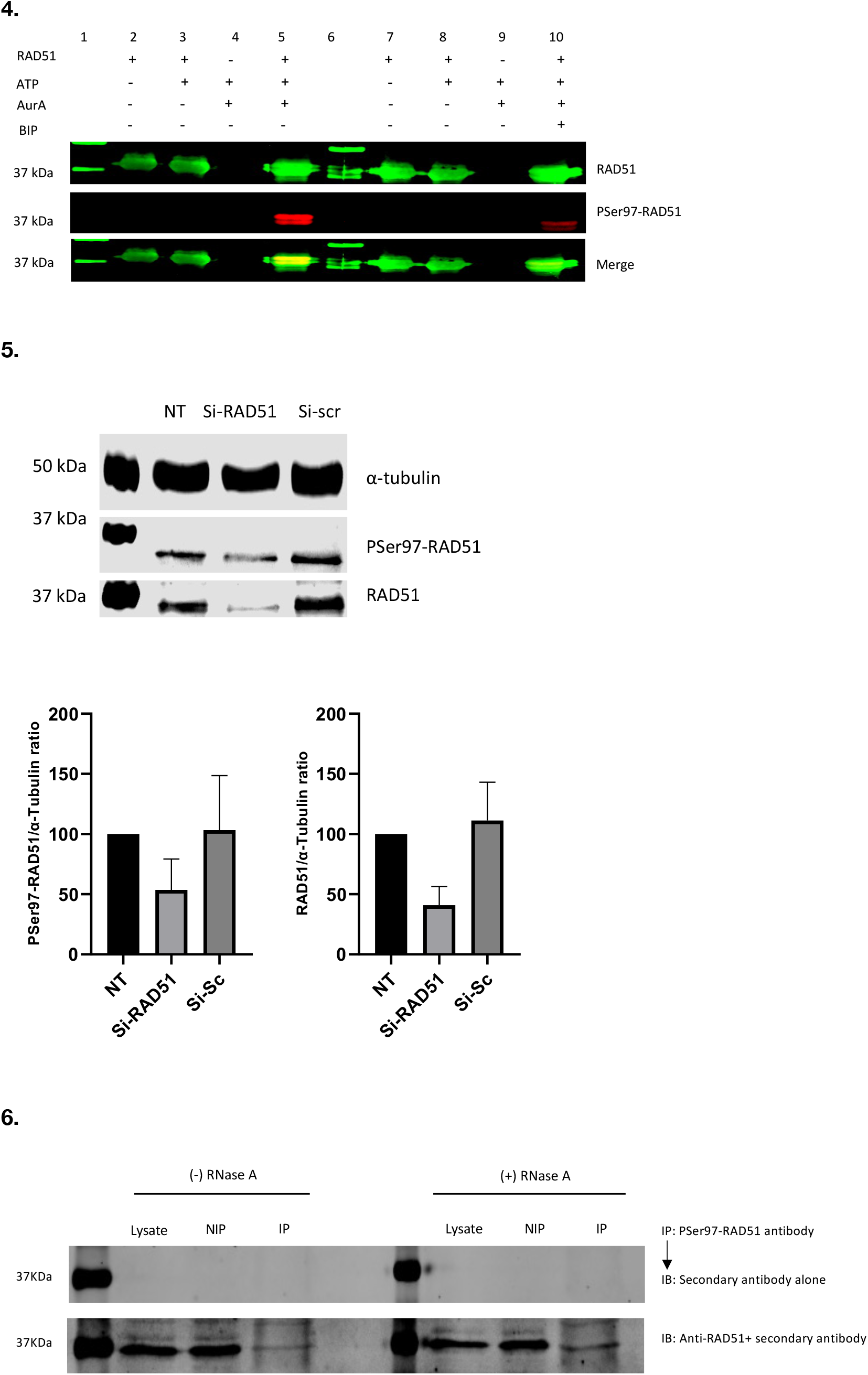
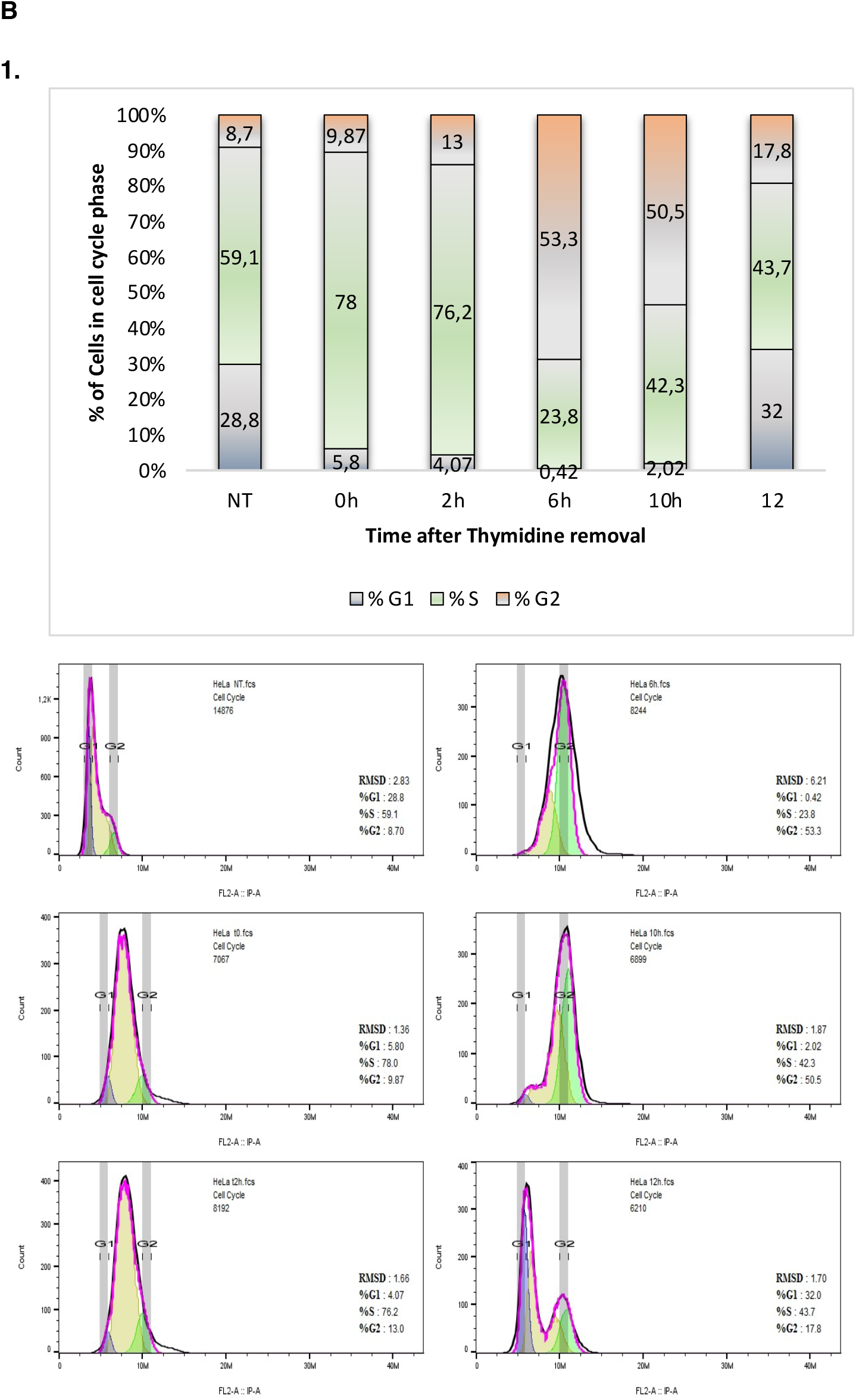
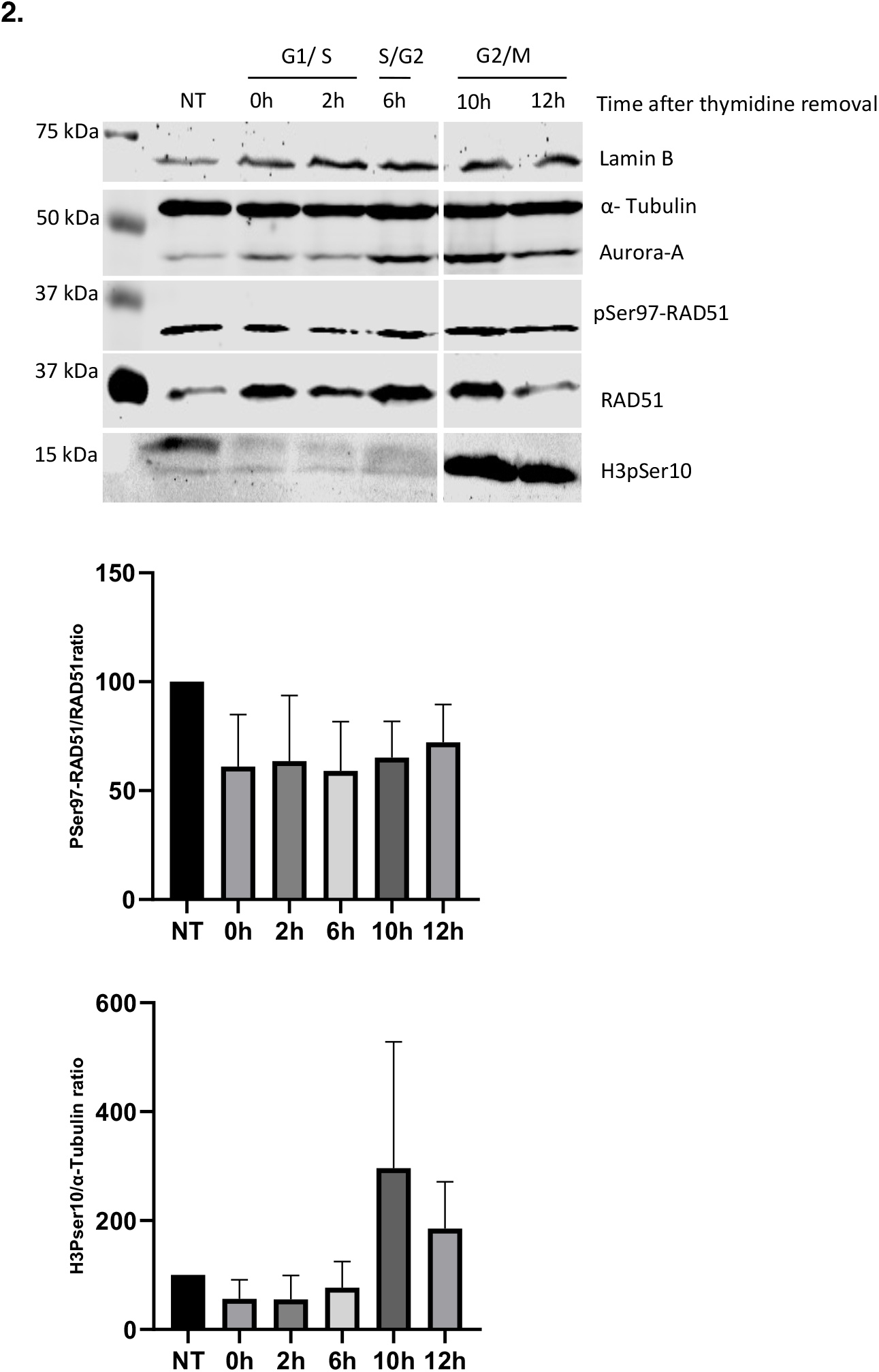
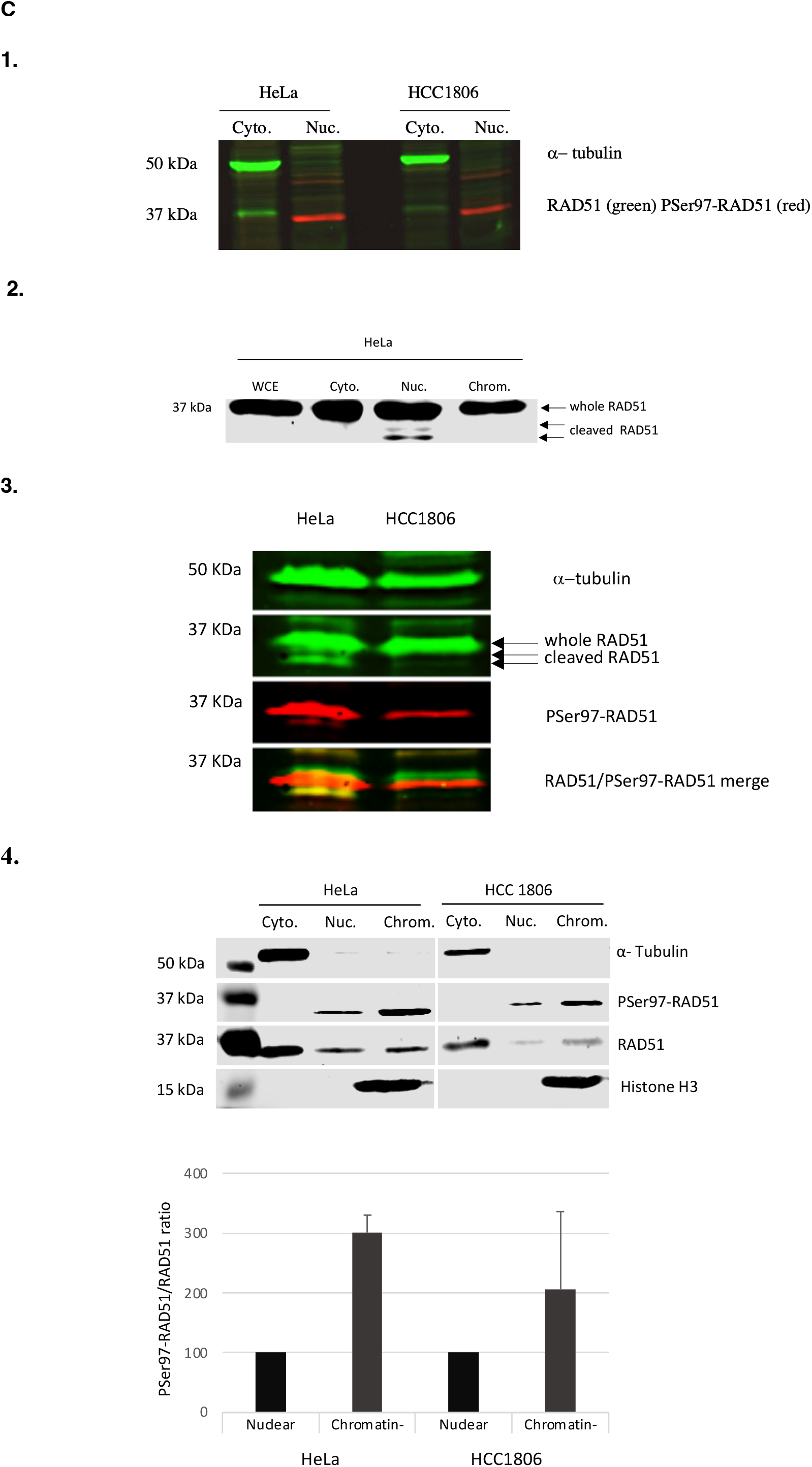
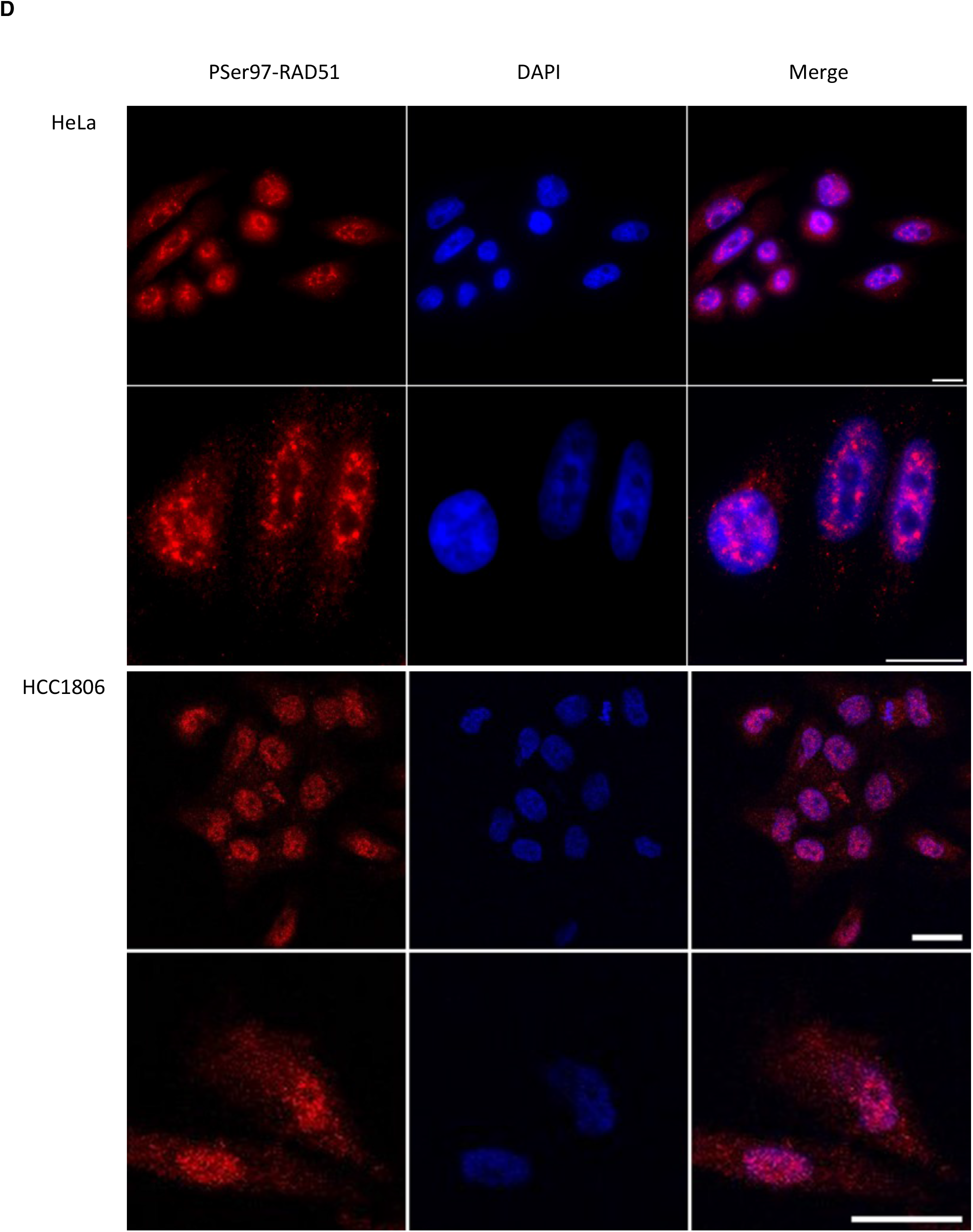
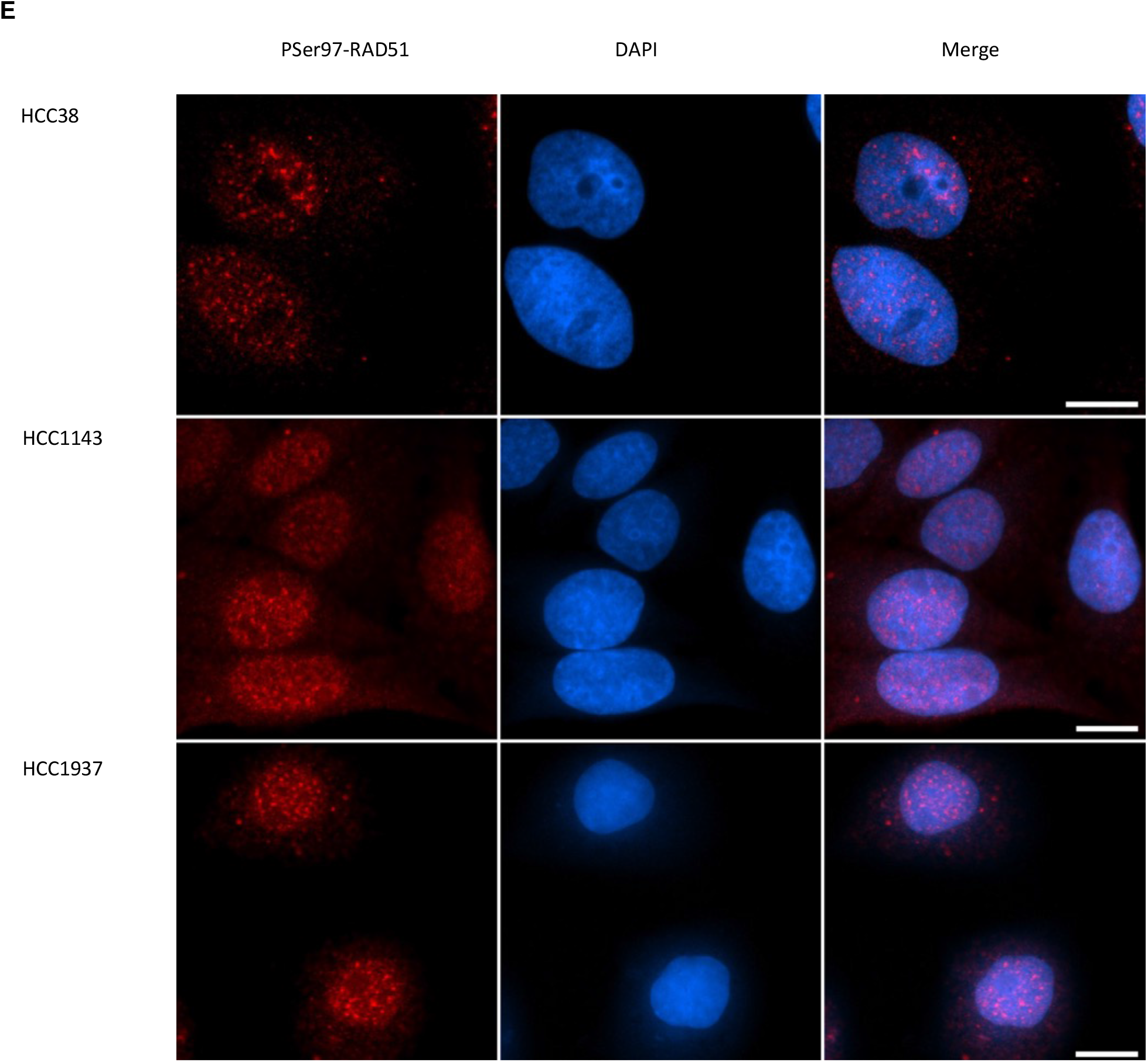
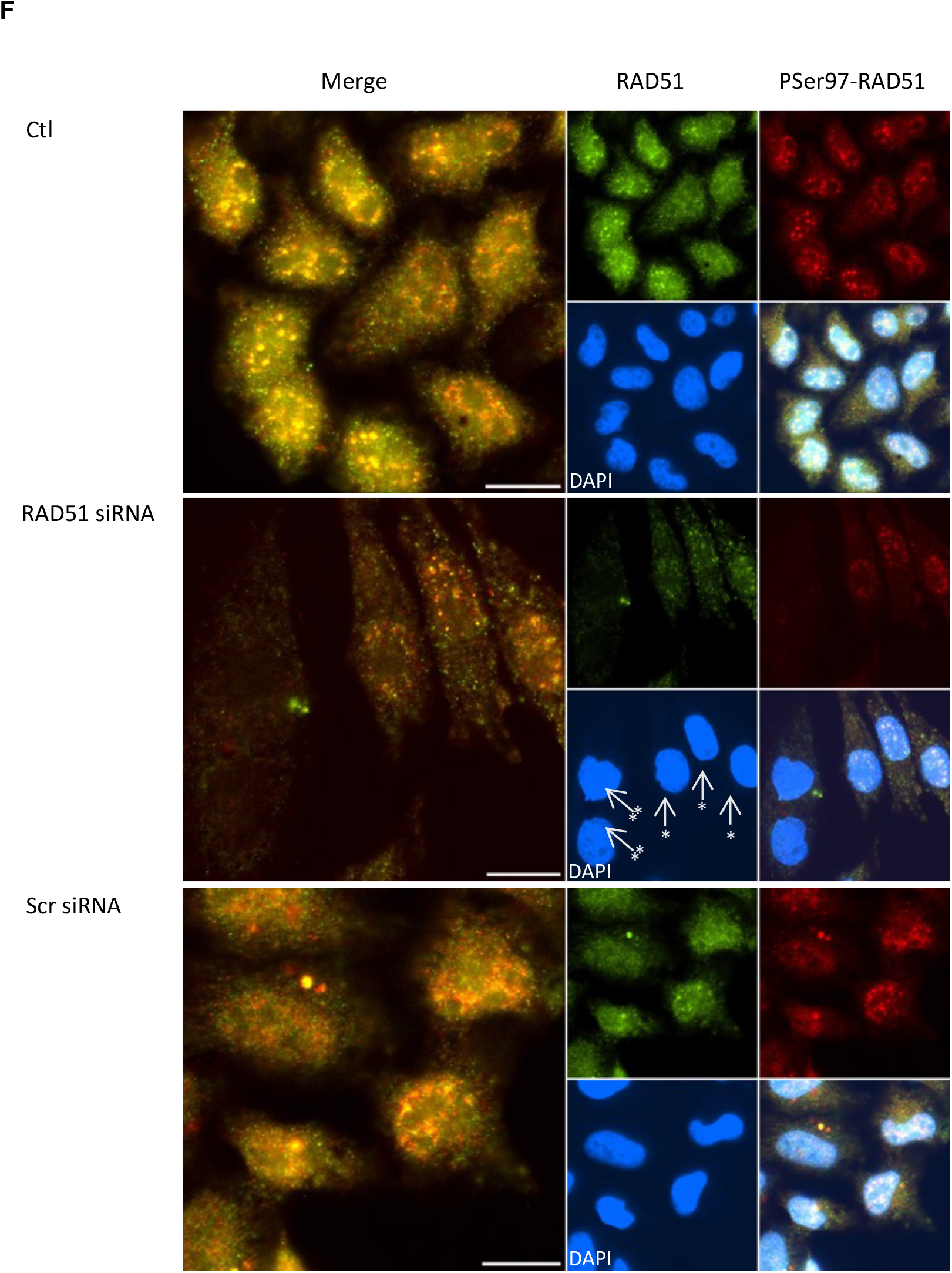
**A: anti PSer-RAD51 antibody validation and *in cellulo* exploration of PSer97-RAD51 presence** 1.Dot blot assay showing the Pser-RAD51 antibody recognition of the PSer-RAD51 peptide and not the Ser-RAD51 peptide. 2. Western blot assay showing the PSer-RAD51 signal in HeLa and HCC1806 whole cell extracts in control condition or after a peptide competition. The dashed lines correspond to the membrane cutting for the two kinds of incubation (classical or with a peptide competition). Two peptides were used for the competition, the non-phosphorylated and the Ser97 phosphorytated one 3. HeLa whole cell extracts were used to perform a western blot in control condition and after the action of Bovine Intestine Phosphatase (BIP). The dashed line corresponds to the membrane cutting for the incubation with BIP of the right part of it. Western blot assay showing an *in vitro* phosphorylation assay of RAD51 by Aurora A kinase in control condition (lines 1-5) and after BIP action (lines 7-10). 5. Western blot assay with HeLa whole cell extracts in control condition and after 24h transfection with RAD51siRNA or scr siRNA. The Pser-RAD51 and RAD51 signals were quantified from n=3 experiments. 6. Immunoprecipitation of PSer97-RAD51 with the phosphospecific antibody, and Immunoblotting using a monoclonal antiRAD51 antibody. RNase A treatment ameliorates the precipitation of the PSer97-RAD51 form. NIP: Non Immuno Precipitated, IP: Immuno Precipitated. **B: PSer97-RAD51/RAD51 ratio is dynamic during the cell cycle progression** 1. Cell cycle progression of HeLa cells after double thymidine block. Cytometer was used to count cells and follow their genomic material after PI (Propidium Iodide) labelling. Cell cycle distribution analysis was performed using FlowJo® software. 2. Western blot assay was used to the tracking of PSer97-RAD51 levels in HeLa cells during cell cycle progression after the double thymidine block. The H3pSer10 was used as a mitosis labelling. **C: PSer97-RAD51 is predominantly localized in the nucleus** Evaluation of the sub-cellular and subnuclear localization of PSer97-RAD51 in control condition in HeLa and HCC1806 cells. 1.Cytoplasmic/nuclear fractionation. Western blot using rabbit PSer97-RAD51 antibody in red and mouse RAD51 antibody in green. Alpha-tubulin was used as a loading control and fractionation quality indicator. 2. HeLa whole cell extracts, cytoplasmic, nuclear soluble and chromatin linked fractions were used to perform a western blot analysis using a commercial anti RAD51 antibody. Undersized RAD51 proteins are observed in the nuclear fraction. 3. The phospho-specific PSer97-RAD51 antibody (red signal) and a commercial antiRAD51 antibody (green signal) were used to check for the presence of this phosphorylation in whole cell extracts of HeLa and HCC1806 cell lines. 4. Western blot assay using PSer97-RAD51 and RAD51 antibodies was performed using cytoplasmic, nuclear soluble and chromatin linked fractions of HeLa and HCC1806 cells. Alpha-tubulin and H3 were used as loading controls and fractionation quality indicators. Signals were quantified and used to calculate the PSer97-RAD51 ratio relative to RAD51 in nuclear soluble and chromatin bound fractions (n=5 experiments). **D: *in cellulo* nuclear enrichment of PSer97-RAD51** Immunofluorescence staining of PSer97-RAD51 in HeLa and HCC1806 cells. PSer97-RAD51 is in red and the DNA DAPI staining is in blue. Scale Bar= 10μm. **E: PSer-RAD51 labelling in different human cell lines** PSer97-RAD51 labelling in HCC38, HCC1143 and HCC1937 breast cancer cell lines. PSer97-RAD51 is in red and the DNA DAPI staining is in blue. Scale Bar= 10μm. **F: *In cellulo* anti PSer-RAD51 antibody validation** HeLa cell line was used to follow the RAD51 and the PSer97-RAD51 signals in control condition and after RAD51-siRNA transfection. Scr siRNA was used as a negative control. In the RAD51 siRNA condition, arrows with one star indicate cells with a diminished RAD51 and PSer97 signals. Arrows with two stars indicate cells with a complete RAD51 and PSer97 RAD51 signal extinction. Scale Bar= 10μm. Image acquisitions properties and treatment were identical for all conditions.

Finally, we performed immunoprecipitation experiments using our specifically generated anti-PSer97-RAD51 antibody raised in rabbits, and a western-blot analysis using a commercial anti-RAD51 antibody raised in mice. Immunoprecipitation of PSer97-RAD51 was difficult to perform and we observed very low levels of immunoprecipitated proteins. We made a first scan of the membrane after its incubation with the secondary antibodies alone, to reveal non-specific signals (cf. upper part of the figure 2A.6). We observe in this figure that RAD51 was revealed after the precipitation with the anti-PSer97-RAD51 antibody and that we slightly improved this experiment by pretreating our whole cell extracts with RNaseA.

All the presented results allowed us to conclude that this generated antibody recognize specifically the Ser97 phosphorylated form of RAD51. This antibody was then used for the *in cellulo* analysis of the PSer97-RAD51using HeLa and HCC1806 cells.

### PSer97-RAD51 is detected at all stages of the cell cycle

HeLa double thymidine-bloc synchronized cells, were used to follow the presence of Ser97-RAD51 phosphorylation by western blotting using whole cell extracts. The cell cycling progression was followed by cytometric analysis of the cells genetic material content and are presented in figure 2B.1. The H3Ser10 phosphorylation was also used to detect mitosis phase. The western blot results are shown in Figure 2B.2. First, we observed that this phosphorylation could be detected throughout the cell cycle. We secondly noticed that the PSer97-RAD51/RAD51 ratio seems to vary a little during the cell cycle progression, but with a statistically non-significant way. We conclude that the PSer97-RAD51 signal is present at all stages of the cell cycle.

### The PSer97-RAD51 is enriched in the nucleus

To evaluate the sub-cellular localization of this new RAD51 phosphorylation, we performed cytoplasmic/nuclear cell extracts fractionations from HeLa and HCC 1806 cell lines. We can see in Figure 2C.1 that the PSer97-RAD51 is observed only in the nuclear fraction of both cells, while the RAD51 signal is mainly cytoplasmic. We also noticed that the PSer97-RAD51 band is below that of RAD51. This point was considered with the recurrent observation made in the lab, of the existence of an undersized nuclear form of RAD51. The figure 2C.2 shows a western blot assay made with a RAD51 directed commercial antibody and illustrates this observation. We can hypothesize that this weaker but recurrent signal corresponds to a cleaved form of RAD51. We then tested the co-incubation of HeLa and HCC1806 whole cell extracts with a RAD51 directed commercial antibody and the PhosphoSer97 antibody. In the figure 2C.3, when observing the results of the HeLa cells, in which the signal is higher, we see clearly that the PSer97 signal is superposed to the undersized RAD51. We concluded that the PSer97-RAD51 is present in the nuclear fraction of HeLa and HCC1806 cell lines.

We then evaluated the subnuclear localization of the PSer97-RAD51 by performing a cytoplasmic/nuclear soluble/chromatin linked fractionation. The results are presented in the figure 2C.4 in which we observed that in the two tested cell lines, the PSer97-RAD51 form is present in both nuclear fractions but is predominantly localized (2 to 3-times more) in the chromatin linked fraction. To validate these results, we used immunofluorescence labelling and confocal microscopy to evaluate the sub-cellular distribution of the PSer97-RAD51 on the same cell lines. The results are presented in Figure 2D. We observed that PSer97-RAD51 (in red) is predominantly localized in the nucleus. The labelling is granular, with a pan-nuclear distribution with nucleolar exclusion and some foci of different sizes that are localized throughout the nucleus. This labeling is shown in figure 2E, for 3 other human cell lines of breast cancers and showed similar results. All these results were obtained under control conditions, meaning in the absence of exogenous DNA damaging agents, and show that the RAD51 Ser97 phosphorylated form is always present in the nucleus. We also used RAD51 directed siRNA to create a RAD51 deficient context in order to validate the anti PSer-RAD51 antibody by immunofluorescence. The results are shown in the figure 2F, in which we observe under RAD51 siRNA condition, the presence of cells without any RAD51 nor PSer97-RAD5 labelling (see the cells labelled with an arrow with two stars). We also observed cells with a diminution of both RAD51 and PSer97-RAD51 labellings (see the cells labelled by an arrow with one star). This experiment validates the use of the PSer97-RAD51 antibody *in cellulo*.

### The phosphorylated Ser97-RAD51 is affected by camptothecin treatment

Camptothecin is an inhibitor of the topoisomerase I that causes replicative stress and DNA damages that are managed by the HR pathway. We tested the effect of a camptothecin-induced DDR on RAD51 Ser97 phosphorylation. We performed western blot experiments using whole cell extracts. We can see in the figure 3A, that the DNA damages induce a decrease of PSer97-RAD51/RAD51 ratio in both cell lines. This decrease was statistically significant only for the HeLa cell line. Therefore, the PSer97-RAD51 decreases after DNA damage induction.

**Figure 3.**
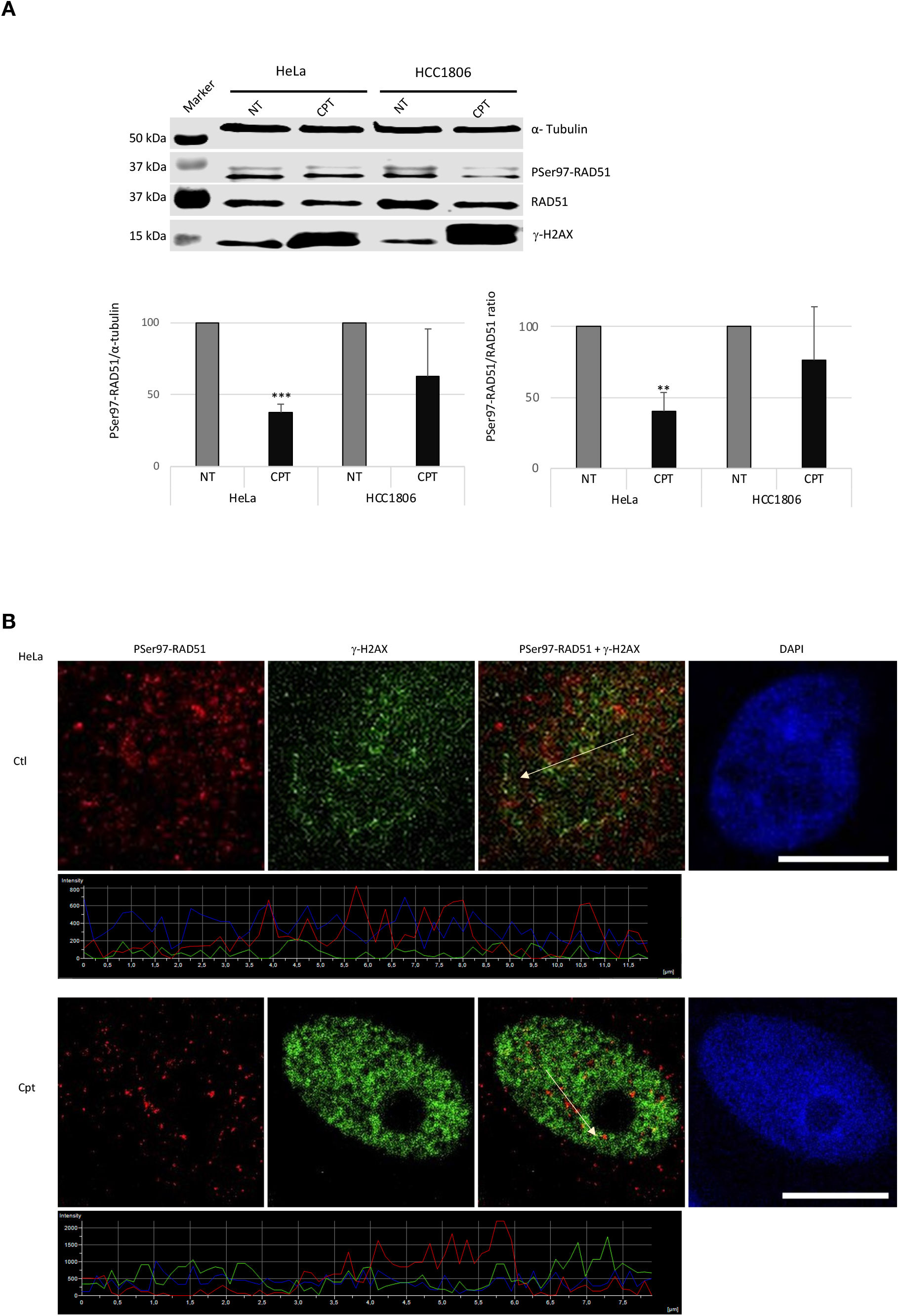
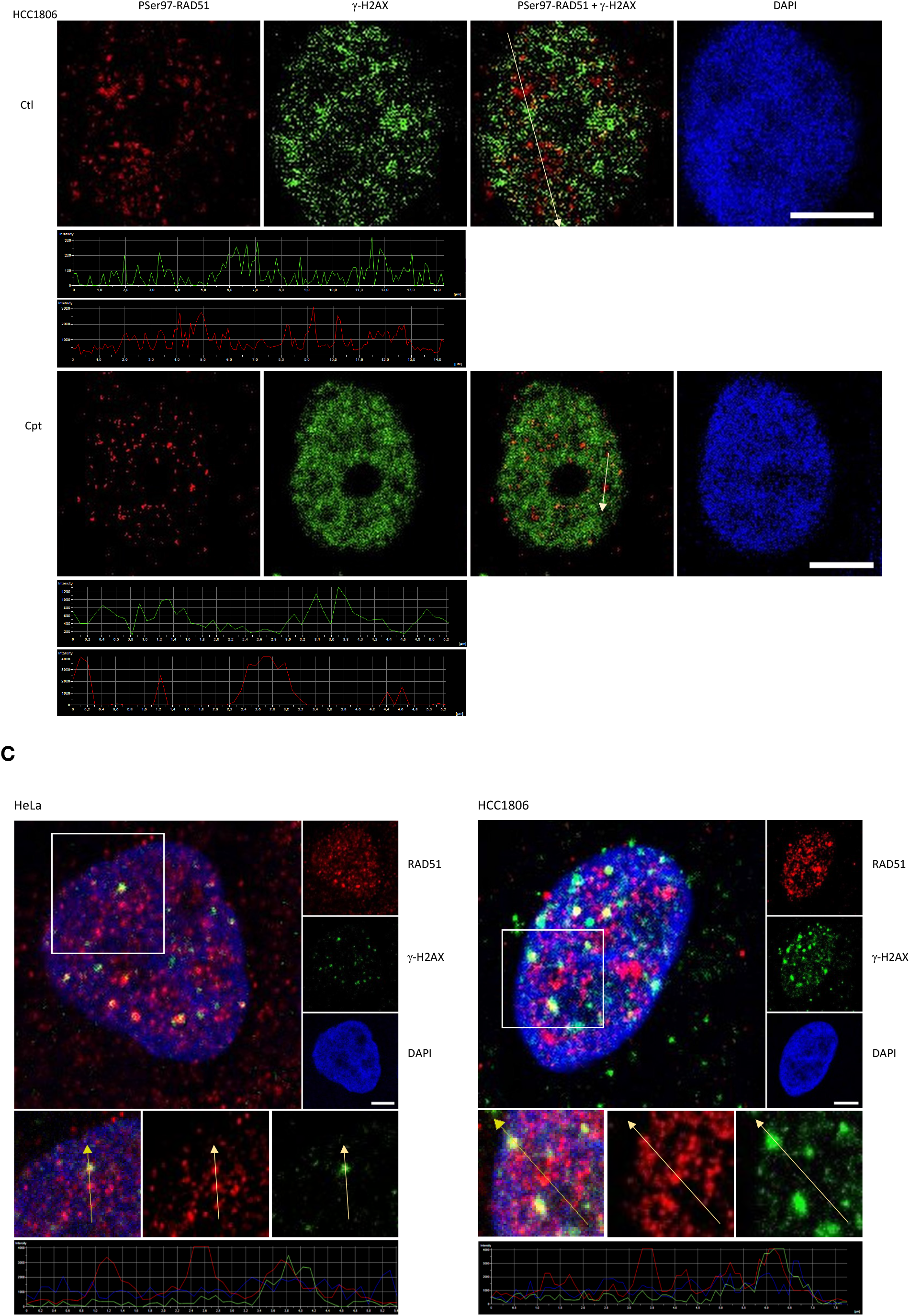
**A: PSer97-RAD51 is reduced after a camptothecin treatment** Evaluation of the Camptothecin-induced DDR effect on PSer97-RAD51 in HeLa and HCC1806 cells. The DDR was induced by a Camptothecin treatment during 4h at 10 μM final concentration. Western blot was performed using HeLa and HCC1806 whole cell extracts, n=4. We observe an enhancement of the γ-H2AX signal after Camptothecin treatment, which induces double strand DNA breaks. PSer97-RAD51/α-tubulin ratio values are: HeLa Ctl: set as 100%, HeLa Cpt= 37.67+/−5.9. t-test p-value (Ctl *vs* Cpt = 0.0014). HCC1806 Ctl: set as 100%, HCC1806 Cpt= 62.6 +/− 32. T-test p-value (Ctl vs Cpt =0.095). PSer97-RAD51/RAD51 ratio values are: HeLa Ctl: set as 100%, HeLa Cpt= 40.5 +/− 13.28. t-test p-value (Ctl *vs* Cpt = 0.008). HCC1806 Ctl: set as 100%, HCC1806 Cpt= 76.5 +/− 37.4 T-test p-value (Ctl vs Cpt =0.19). **B: PSer97-RAD51 foci are not localized in DNA damaged sites** HeLa and HCC1806 cells in control condition and after Camptothecin treatment (10 μM for 4h) conditions were used for immunofluorescence labelling of PSer97-RAD51. PSer97-RAD51 is in red, γ-H2AX is in green and the DNA DAPI staining is in blue. Line scan analysis showing the PSer97-RAD51 and γ-H2AX signals intensities along the arrow are below each image. Scale Bar= 10μm. **C: Some RAD51 foci are not DD sites** Confocal images of immunofluorescence labelling of RAD51 and γ-H2AX with commercial antibodies in non treated HeLa and HCC1806 cells. RAD51 is in red, γ-H2AX is in green and the DNA DAPI staining is in blue. Scale Bar= 10μm. The squares indicate magnified areas used for line scan analysis of the signal intensities along the arrows.

In this context, we performed immunofluorescence experiments to evaluate the effect of camptothecin on PSer97-RAD51 sub-cellular localization. Here again, we labelled the PSer97 in red and the γ-H2AX DNA damages foci in green. For reminder, the DNA double strand breaks are classically marked by the presence of a phosphorylated form of the histone H2AX variant, called γ-H2AX which is found enriched on several kb around the DNA breaks ^25^. The results are shown in figure 3B. We were very surprised to observe that the PSer97 foci were not colocalized with the γ-H2AX foci. This was true for the control condition, in which we observed dsDNA breaks due to the replicative stress, and the camptothécine treatment condition. We used confocal images and a line scan analysis to illustrate that observation. We can see in the figure 3B that the two labelings are completely different, meaning that PSer97-RAD51 is not located in DNA damage “repair centers”. To validate these atypical results, we performed a labelling of RAD51 using a commercial antibody and checked if there were some RAD51 foci that do not colocalize with γ-H2AX foci. The results are shown in Figure 3C and we can see that there are indeed some RAD51 foci that are not colocalized with γ-H2AX foci. Here again, we used confocal microscopy and line scan analysis tool to illustrate this observation. We easily see in the figure 3C that some RAD51 foci are not colocalized to γ-H2AX foci.

### Aurora A kinase is implicated in the phosphorylation of RAD51 on its Ser97 residue

We identified the Ser97-RAD51 phosphorylation *in vitro*, under the action of the Aurora A kinase, but the complexity of the *in cellulo* mechanisms regulating protein phosphorylation and the multiple redundancy observed between kinases, do not allow us to consider that Aurora A kinase is necessarily also responsible for this phosphorylation *in cellulo.* We explored the implication of the Aurora A kinase in the *in cellulo* phosphorylation of RAD51 by using two approaches to enhance or diminish Aurora A kinase activity. On the one hand we used alisertib, a known specific Aurora A inhibitor, and on the other hand we performed an Aurora A over-expression, and we assessed the effect of these two conditions on RAD51-Ser97 phosphorylation. The results of Aurora A inhibition are shown in Figure 4A, in which we see the PSer97RAD51/RAD51 ratio for all conditions. The Aurora A inhibition induced a statistically significant decrease of PSer97-RAD51 in the HCC1806 cell line, while it had no effect on the HeLa cells.

**Figure 4:**
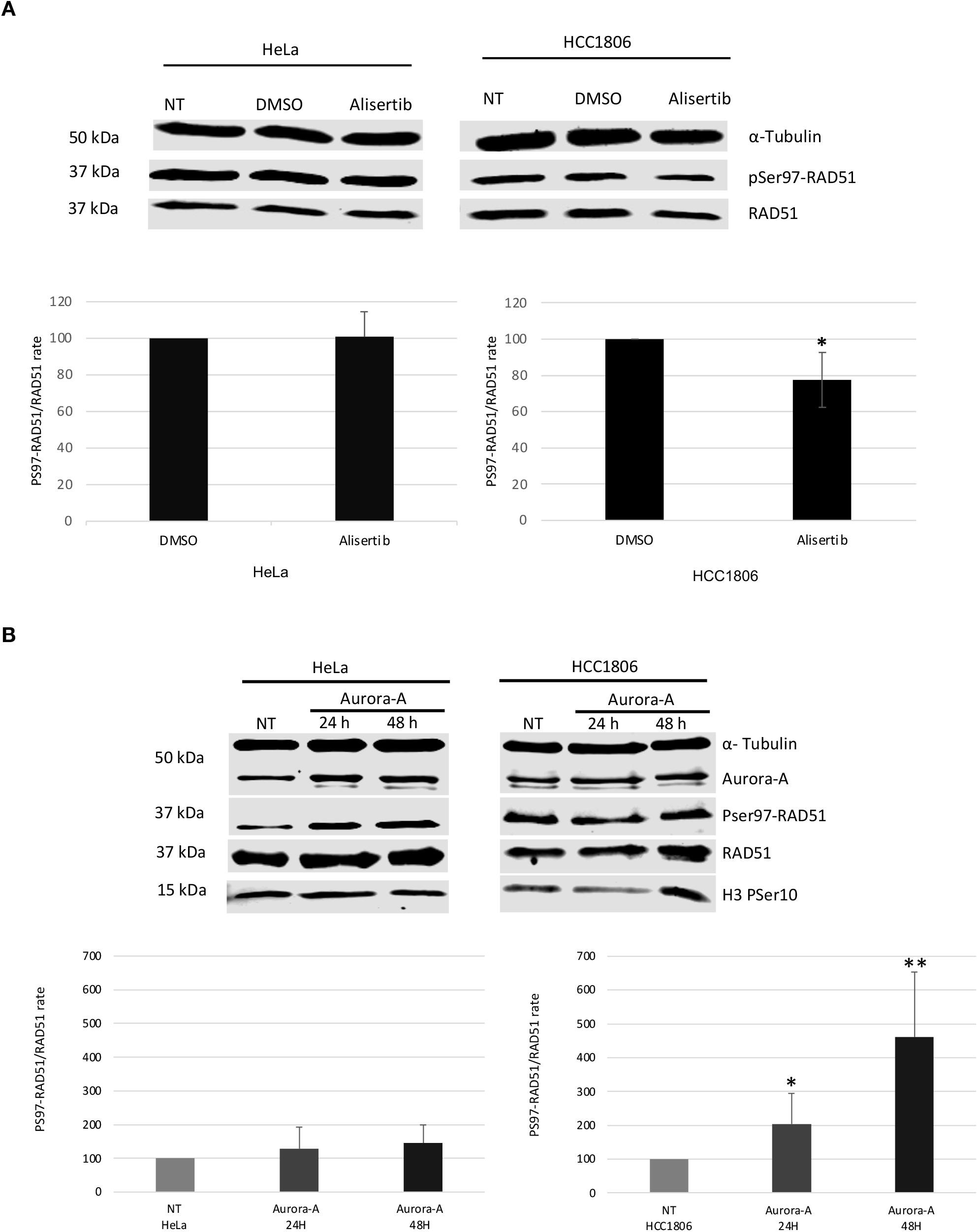
Aurora A inhibition/ overexpression affect PSer97-RAD51 cellular level. **A.** Evaluation of the use of Aurora A inhibitor on PSer97-RAD51. The Aurora inhibitor Alisertib was used at a 50nM final concentration for 24 hours in HeLa and HCC1806 cells. Statistical analysis was done for n=5 experiments, using a paired t-test. p-value (α= 0,05: *, α= 0,001: **, α= 0,001: ***). PSer97-RAD51/RAD51 ratio values: HeLa DMSO: set as 100%, HeLa Ali =100.7 +/− 13.7, (p-value DMSO *vs* Ali = 0.9). HCC1806 DMSO: set as 100%, HCC1806 Ali= 77.7 +/− 15.18, (p-value DMSO *vs* Ali = 0.03). **B.** Evaluation of the effect of Aurora A over-expression on PSer97-RAD51. Aurora A coding plasmid was transfected for 24h and 48 h in HeLa and HCC1806 cells. WB were done with whole cell extracts and the PSer97RAD51/RAD51 ratio was calculated. Statistical analysis (using a paired t-test) was done for n=3 experiments, p-value (α= 0,05: *, α= 0,001: **, α= 0,001: ***). PSer97-RAD51/RAD51 ratio values: HeLa NT: set as 100%, HeLa 24h =128.1 +/− 64, (p-value NT *vs* 24h = 0.2), HeLa 48h =145 +/− 54.4, (p-value NT *vs* 48h= 0.09). HCC1806 NT: set as 100%, HCC1806 24h = 203 +/− 90, (p-value NT *vs* 24h = 0.050), HCC1806 48h = 461+/− 191, (p-value NT *vs* 48h = 0.016).

Aurora A over-expression by transient transfection was performed and the total extracts were used to evaluate the PSer97-RAD51/RAD51 ratio. In figure 4B, we observe that Aurora A over-expression induced an enhancement of the PSer97-RAD51/RAD51 ratio in both cell lines, and the statistical analysis allowed us to conclude that this enhancement is significant in the HCC1806 cell line. Therefore, the aurora A inhibition or overexpression experiments performed in the HCC1806 cell line allowed us to conclude that Aurora A is implicated in the *in cellulo* phosphorylation of RAD51 on its Ser97 residue. The absence of a statistically significant effect in the HeLa cell line underlines the probability of a complex regulation of RAD51 Ser97 phosphorylation in this cell line. This point will be developed in the discussion part.

### The phosphorylated Ser97-RAD51 is located in the Nuclear Speckles

To identify the structures in which we observe the PSer97-RAD51 foci, we performed immunofluorescence experiments and labelled different nuclear organelles. Nuclear Speckles (NS) are membrane less organelles, also called interchromatin granules, which contain RNA, RNA binding proteins (RBP) and splicing factors. These structures are the location of alternative splicing and mRNA maturation. Using immunofluorescence co-labelling and confocal microscopy, we showed that the PSer97-RAD51 foci are colocalized with Sc35, a Nuclear Speckles component.

We show in the figure 5A a colocalization analysis of the Sc35 and the PSer97-RAD51 labelling on confocal images. The line scan analysis along the arrow shows that maximal pics intensities of the two signals are perfectly colocalized. We used confocal microscopy acquired images to quantify the colocalization of the Sc35 and PSer97-RAD51 signals using JACoP tool on Fiji. We used the Pearson’s Correlation Coefficient (PCC) analysis that is classically used to address this type of question. For reminder, the PCC value varies from −1: which means exclusion of the two signals, to +1: which means a total colocalization of the signals. A PCC equal to 0 means a random distribution of the two signals. According to the interpretations standards detailed in the material section, PCC values were used to interpret more finely the colocalization ^23^. We obtained a PCC=0.608+/− 0.05 for HeLa cells and a PCC=0.555+/− 0.04 for HCC1806 cells. These results allowed us to conclude to a high colocalization of these two signals in the two cell lines, which signifies the colocalization of the PSer97-RAD51 within NS. To answer if this RAD51 localization within the Nuclear Speckles is linked or not to a splicing activity, we performed experiments in which we inhibited the pre-mRNA maturation. Pladienolide B (plaB) is a splicing-inhibitor that interacts with the SF3b1 subunit of the spliceosome. This drug leads to the accumulation of incompletely spliced or unspliced pre-mRNA, and provokes modifications of the Nuclear Speckles that become larger in size with a modified shape ^26^. We treated HeLa and HCC1806 cells with plaB to evaluate the effect on the PSer97-RAD51 foci.

**Figure 5:**
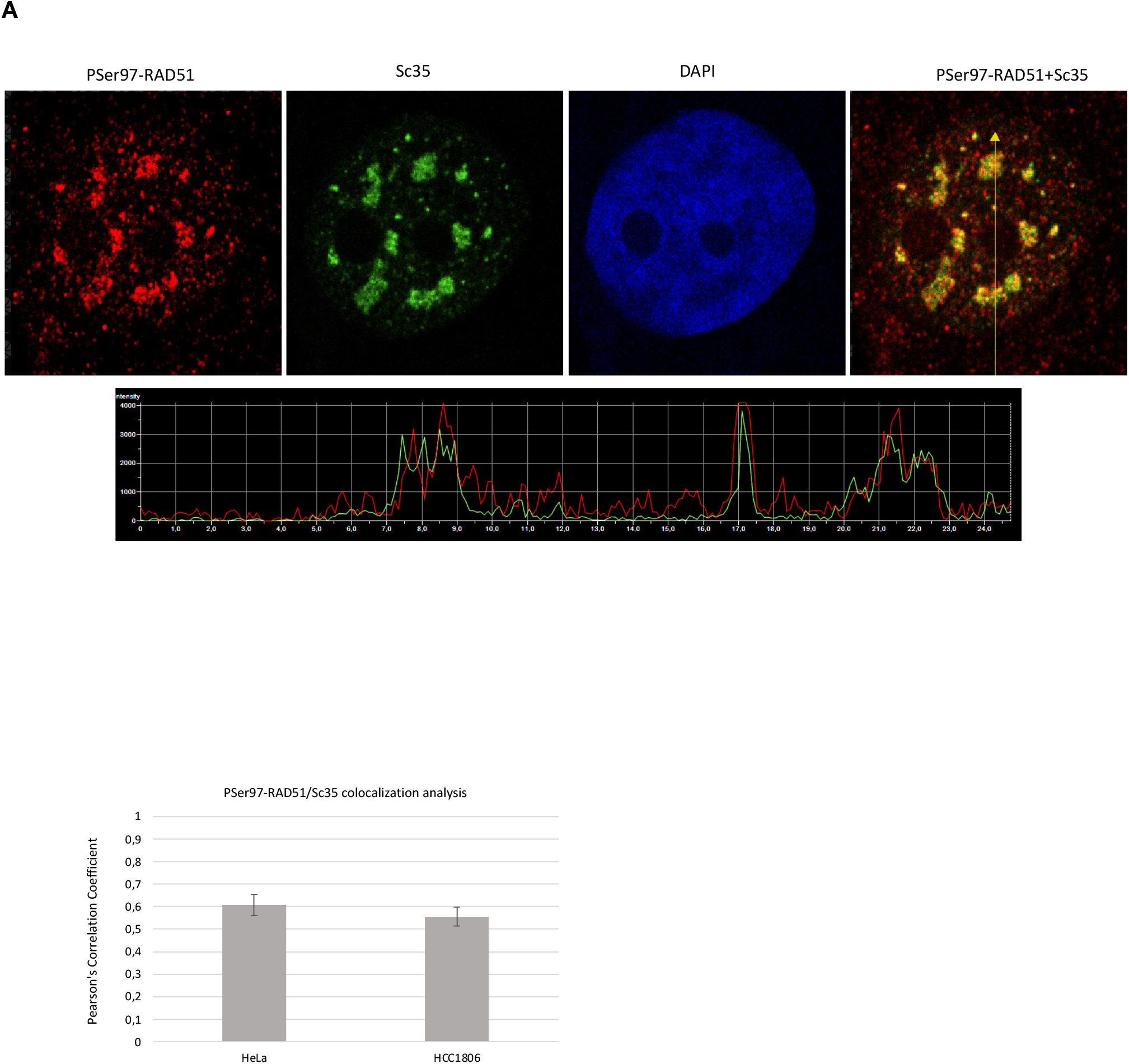
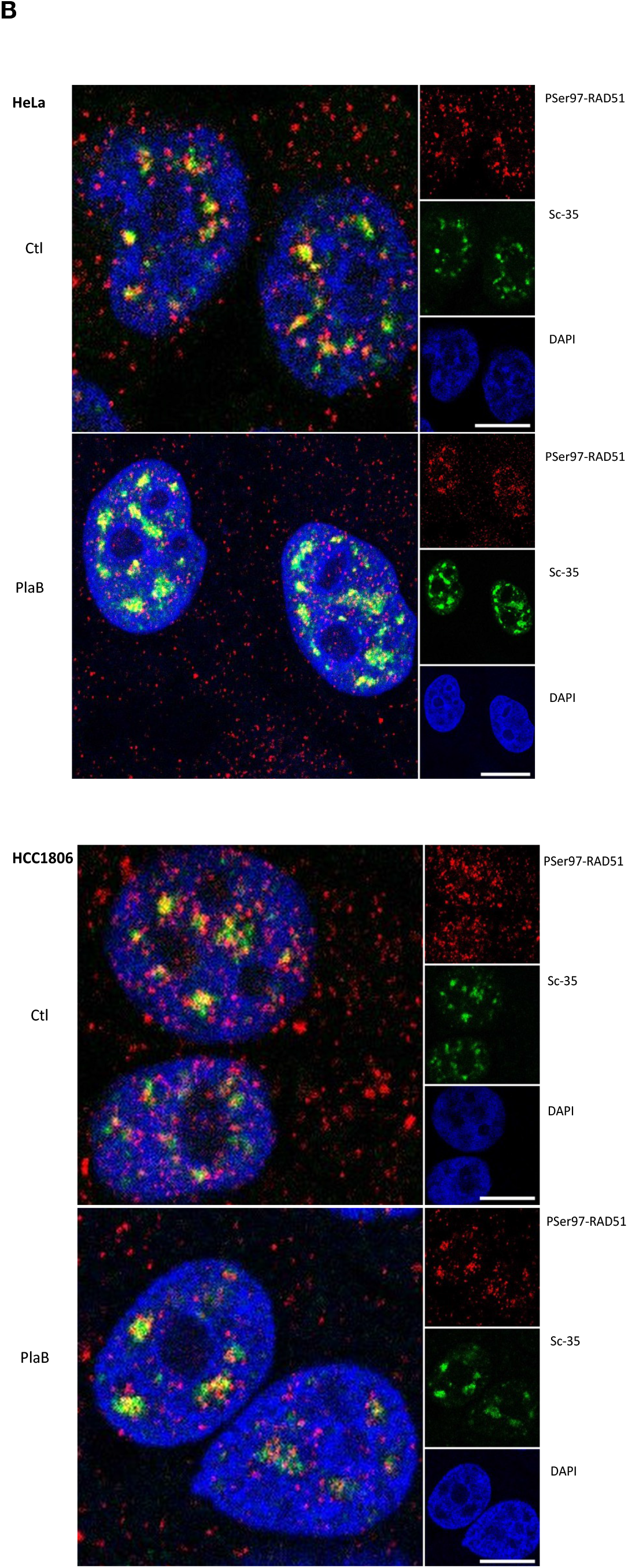
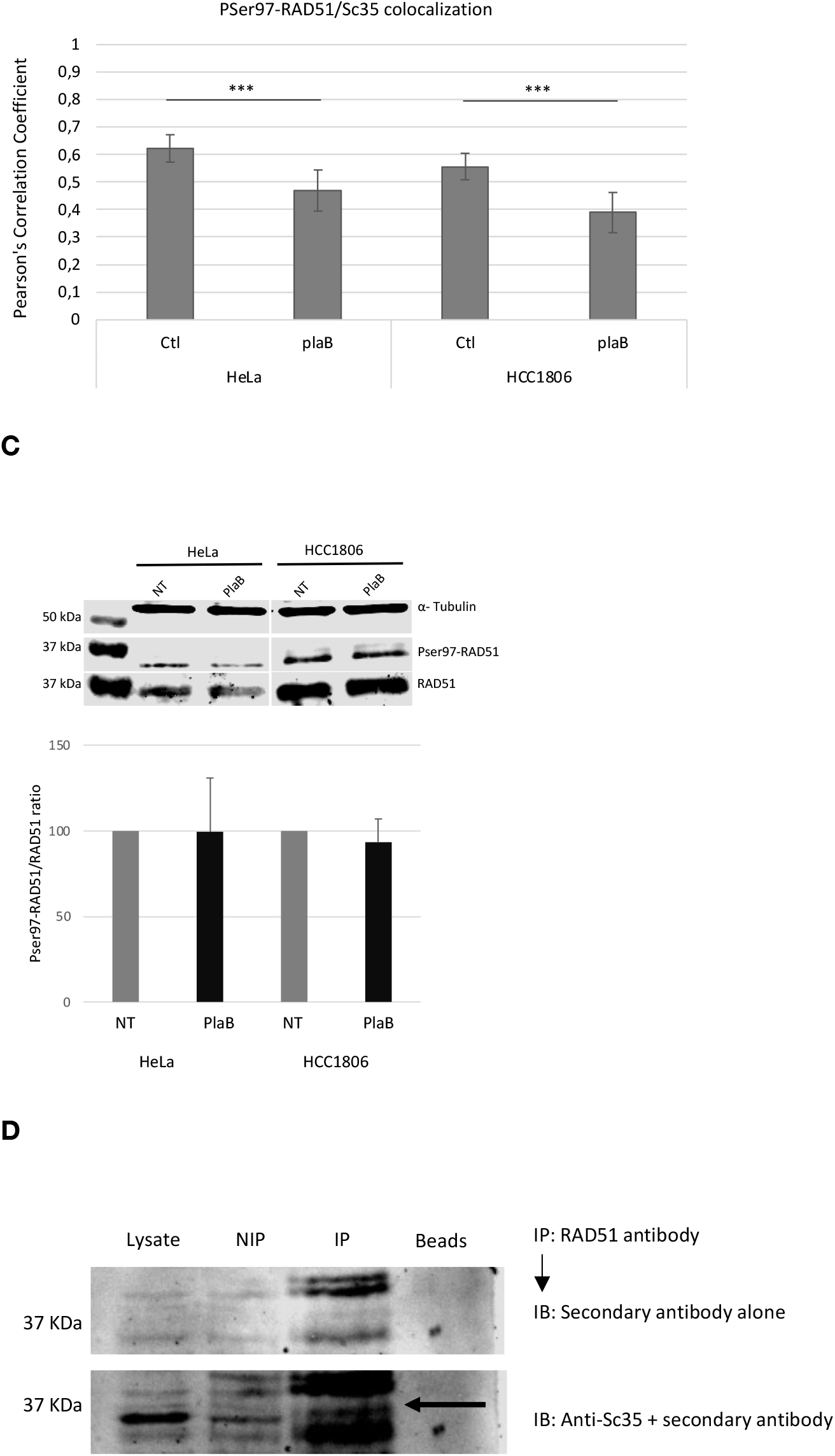
PSer97-RAD51 nuclear foci are localized within Nuclear Speckles and remain there after pladienolide B treatment. **A.** Line scan analysis showing the PSer97-RAD51 and Sc35 signal intensities along the arrow, (HeLa cells). PSer97-RAD51/Sc35 colocalization was evaluated by a Pearson’s Correlation Coefficient (PCC) analysis using confocal images and JACoP/Fiji. For HeLa: PCC= 0,608 +/− 0,05, n= 30 and for HCC1806, PCC= 0,555 +/− 0,04, n=35 cells. PCC: Pearson’s Correlation Coefficient. Results are mean +/− sd (sd: standard deviation) **B.** Immunofluorescence staining of PSer97-RAD51 in HeLa and HCC 1806 cells. PSer97-RAD51 is in red, the nuclear speckles component Sc35 is in green and the DNA DAPI staining is in blue. Confocal acquisitions. Scale Bar= 10μm. PlaB effect on PSer97-RAD51/Sc35 colocalization was evaluated by a Pearson’s Colocalization Coefficient (PCC) analysis using confocal images and JACoP/Fiji. For HeLa ctl: PCC= 0,608 +/− 0,05, n= 30. For HeLa plaB: PCC=0.468 +/− 0.06, n=32. For HCC1806 ctl, PCC= 0,555 +/− 0,04, n=35. For HCC1806 plaB: PCC= 0.388 +/− 0.07, n=28. One tailed t-test was used to evaluate plaB effect on PCC values. For HeLa p-value= 1.17E-6 and for HCC1806 p-value= 5.04E-7. Results are mean +/− sd **C.** PSer97-RAD51 level is not affected by plaB treatment. HeLa and HCC1806 cells were treated by pladienolide B at 5 μM, 4h. Whole cell extracts were used to evaluate the effect on PSer97-RAD51/RAD51 ratio by Western-Blot analysis. PSer97-RAD51/RAD51 values are: HeLa ctl : set as 100%, HeLa plaB= 99,4 +/− 31.7, HCC1806 Ctl: set as 100%, HCC1806 plaB= 93.3 +/− 13.5. Results are mean +/− sd. **D.** Sc35 is co-immunoprecipitated with RAD51 from HeLa nuclear extracts. NIP: Non Immuno Precipitated, IP: Immuno Precipitated. The black arrow indicates the Sc35 signal.

We can see in the figure 5B that the PSer97 signal is still present with large foci which remain co-localized with the Nuclear Speckles after plaB treatment. Here again, we quantified the colocalization of PSer97-RAD51 and Sc35 in control and plaB conditions, using the PCC indicator. Results are shown in the figure 5B. We found that the PCC values after plaB treatment were still typical to a colocalization of the two analyzed labelings. However, we noticed that the plaB treatment induces a statistically significant decrease of theses PCC values. For the HeLa cell line, the PSer97-RAD51/SC35 colocalization is strong in control condition (PCC= 0,608+/−0.05) and moderate after plaB treatment (PCC= 0.468+/− 0.06). For HCC1806 cell line too, the colocalization is strong in the control condition (PCC= 0.555+/− 0.04) and moderate after plaB treatment (PCC= 0.388+/− 0.07). Meaning that PSer97-RAD51 stays colocalized within NS even when splicing is inhibited.

To evaluate the impact of this treatment on the PSer97-RAD51 level we used western-blot analysis and signal quantifications. The results presented in the figure 5C show that there was no statistically significant difference after plaB treatment. Meaning that the PSer97-RAD51 signal quantity at the cellular level is not affected by plaB induced splicing inhibition.

In order to evaluate the *in cellulo* interaction between RAD51 and Sc35, we performed immunoprecipitation assays of HeLa nuclear extracts using a commercial anti-RAD51 antibody. The western blot membrane was scanned a first time just after its incubation with the secondary antibody alone. The obtained signals are non-specific and are shown in the upper part of the figure 5D. After that, a classical immunoblot was performed to reveal Sc35 factor. The results in the lower part of the figure 5D and show (at the level of the arrow) that Sc35 factor is co-immunoprecipitated with RAD51. These results confirm the immunofluorescence experiments showing that RAD51 exits in the same structure as Sc35 within Nuclear Speckles.

### RAD51 overexpression affects the Nuclear Speckles

To pursue our investigation of the presence of RAD51 within the Nuclear Speckles, we tested the effect of RAD51 overexpression on these structures. We used plasmids coding the HA-tagged RAD51 protein, either Wild Type (WT) or mutated on the Ser97 residue, (S97A non phosphorylable mutant, and S97D phosphomimetic mutant). HeLa and HCC1806 cell lines were transfected with these plasmids for 24h before their use for immunofluorescence labelling of the HA-tag or Sc35 epitopes. The results obtained with the HeLa and HCC1806 cells are shown in figure 6 panels A and B respectively. In the HeLa cells, we observed that exogenous WT and mutated RAD51 were translocated into the nucleus. RAD51 overexpression and its nuclear translocation had a strong effect on the Nuclear Speckles. We can see that the transfected cells, visualized by the HA-labelling and white arrows, had an abnormal Sc35 labeling. Indeed, in these transfected cells, the Nuclear Speckles number were strongly diminished, and some cells had no NS at all. This effect was observed for the WT as well as for the two mutants. The same experiment was performed in the HCC1806 cells, in which we observed that the exogenous WT and mutant RAD51 were expressed, but maintained in the cytoplasm. Besides this difference in exogenous RAD51 sub-cellular localization, we observed that RAD51 overexpression in HCC1806 cells had no “eye visible” strong effect on NS and we observed no cell without NS. Using Image J, we quantified the NS number for the different conditions in order to determine the average NS number per cell. The results are presented in the figure 6C. In HeLa cells, consistently with our observations, after the overexpression of exogenous RAD51, the average NS number/cell is strongly diminished in a statistically highly significant manner, with the presence of cells without NS. Concerning the Ser97 residue, we noticed that the S97A mutant has the lower average NS number per cell, while the S97D and WT RAD51 gave the same phenotype. Surprisingly, in the HCC1806 cell line, the NS quantification revealed that the WT and S97D overexpression had also a diminution of the average NS number /cell that is statistically significant. In this same cell line, the S97A overexpression had no effect at all.

**Figure 6:**
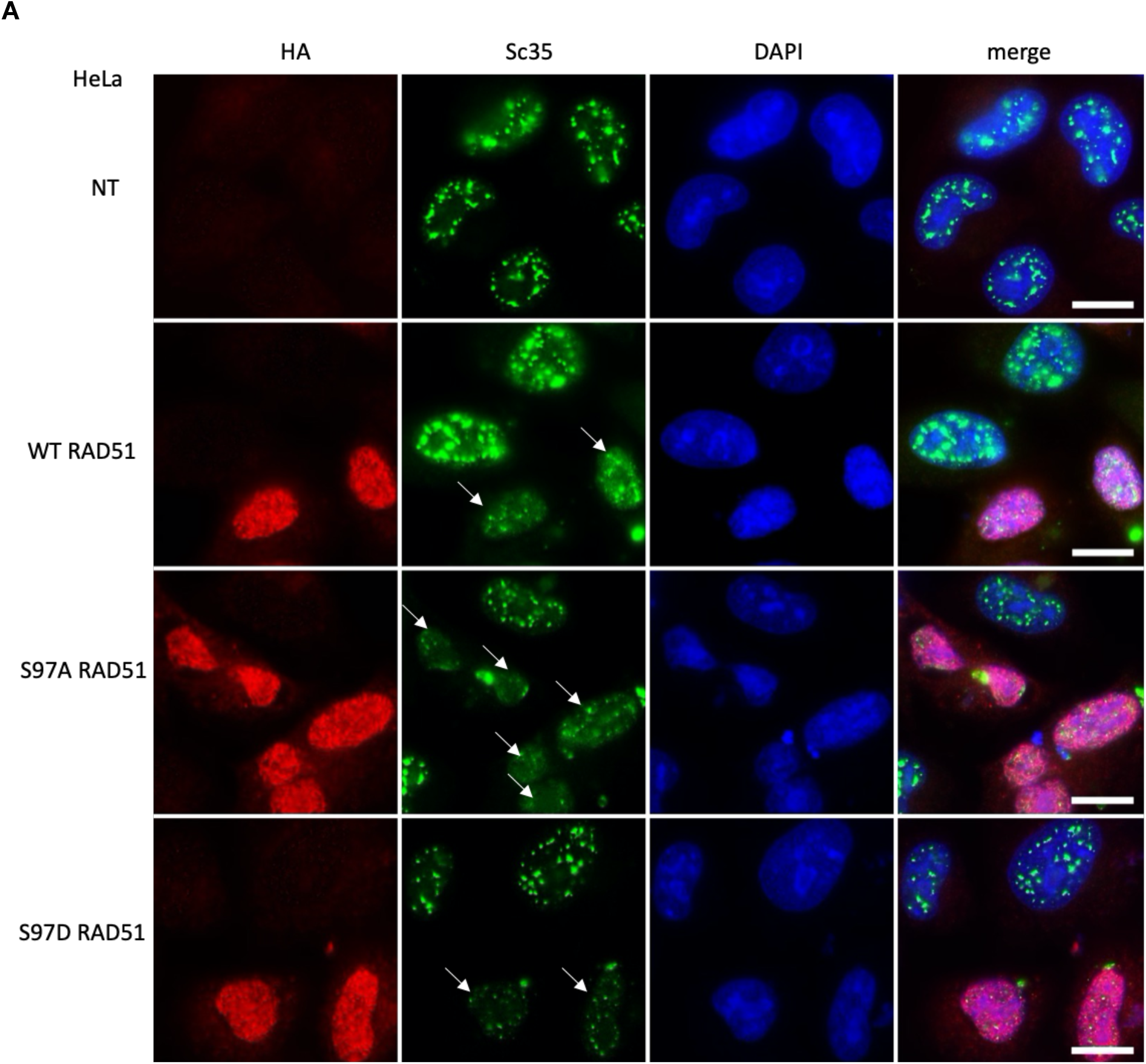
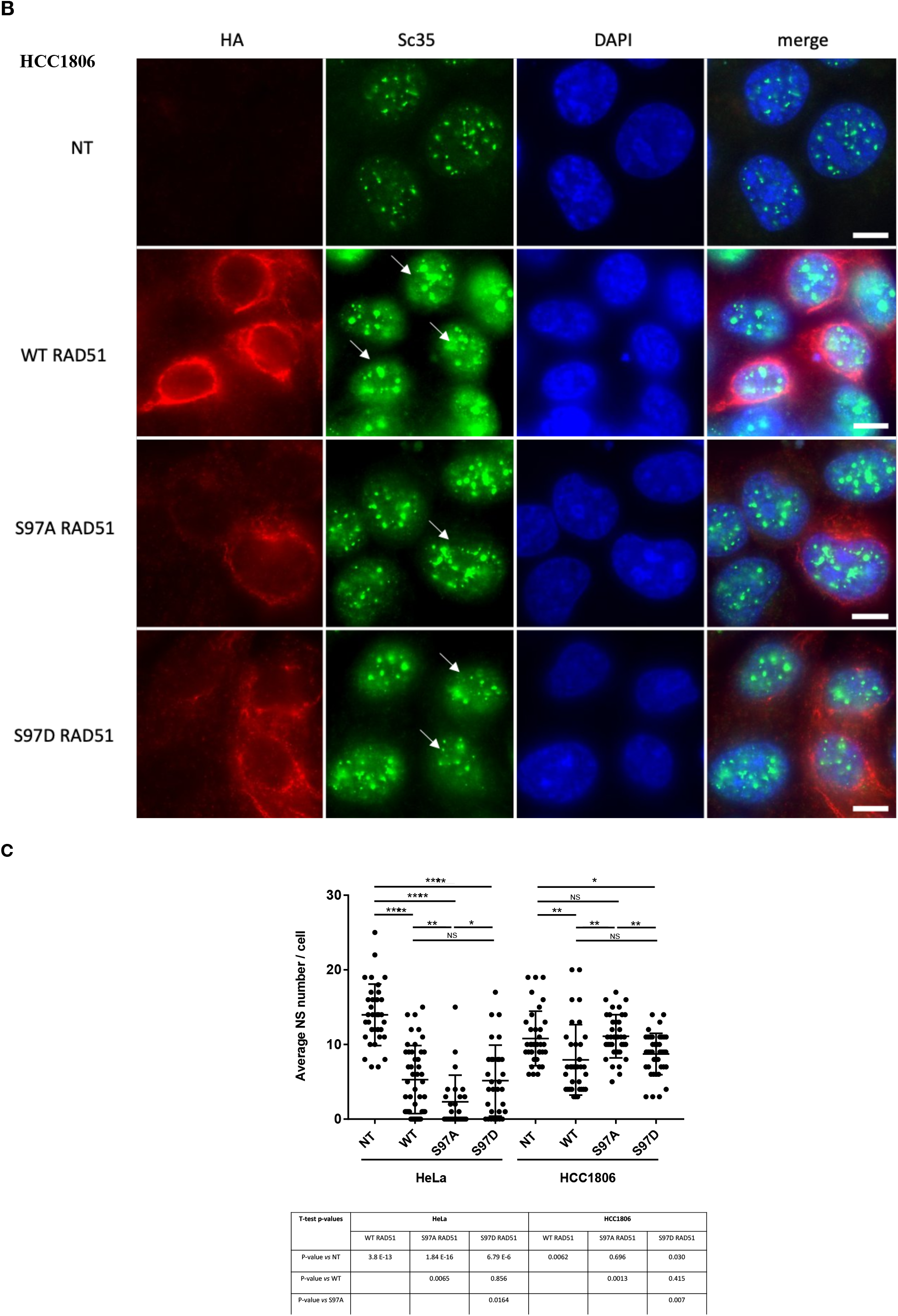
RAD51 overexpression affects Nuclear Speckles number. **A, B:** HA tagged WT, S97A and S97D RAD51 proteins were expressed during 24h in HeLa (panel A) and HCC1806 (panel B) cells. Exogenous RAD51, in red, was labelled with the anti-HA antibody. NT: Non-Treated. The arrows indicate the RAD51 overexpressing cells. The nuclear speckles component Sc35 is in green and the DNA DAPI staining is in blue. Epifluorescence microscope acquisitions. Scale bar= 10 μm. **C**: Average Nuclear Speckles number per cell was quantified using FiJi analysis tool with confocal images. (n=32 to 51 cells for Non-Treated (NT)and 24h transiently transfected HeLa and HCC1806 cells. Two tailed t-tests were performed to evaluate the statistical significance of the observed differences after WT, S97A and S97D RAD51 overexpression on average NS number/cell. Table of the obtained p-values.

Taken together, these results showed that exogenous RAD51 overexpression induces a decrease of the average Nuclear Speckles number per cell, in the two cell lines, but with a more pronounced way in the HeLa cells, where exogenous RAD51 is localized within the nucleus. The most striking difference between the two cell lines concerns the presence of cells without NS. A phenotype that is observed only in HeLa cells and particularly those overexpressing the S97A non phosphorylable RAD51 mutant. In comparison, within the HCC1806 cell line, the S97A overexpression maintained within the cytoplasm had no effect on average NS number per cell. Thus, nuclear translocation of the exogenous non phosphorylable RAD51 is an essential point associated with a NS destabilization.

### RAD51 is an RNA binding protein and its Ser97 phosphorylation affects its binding to RNA

After the finding of RAD51 colocalization within nuclear structures highly enriched in RNA, we pursued our investigations by testing RAD51 ability to bind RNA. In the context of DNA damage repair *via* the HR pathway, RAD51 is a well-known DNA binding protein, that has also been described as promoting DNA/RNA hybrids structures called R-loops, with the TERRA long non coding RNA in the context of telomeric maintenance ^27^. In this same work, they showed that RAD51 binds the TERRA RNA *in vitro*. Little is known about RAD51 affinity to bind RNA. Using Blitz technology, we performed *in vitro* experiments to compare RAD51 binding to ssDNA *vs* ssRNA and evaluate the impact of the phosphorylation on the Ser97 residue on these bindings. For this, we used the phosphomimetic and non phosphorylable recombinant proteins described in the materials section. The results are presented in the figure 7A and B, in which association and dissociation curves are presented for 33 nt sized DNA and RNA respectively. Different RAD51 concentrations were used in order to determine the equilibrium dissociation constant KD of the WT and mutant RAD51 proteins for DNA and RNA binding. The curves are presented in the fig 7C for DNA and fig 7D for RNA, the estimated KD values are presented in the figure 7E.

**Figure 7:**
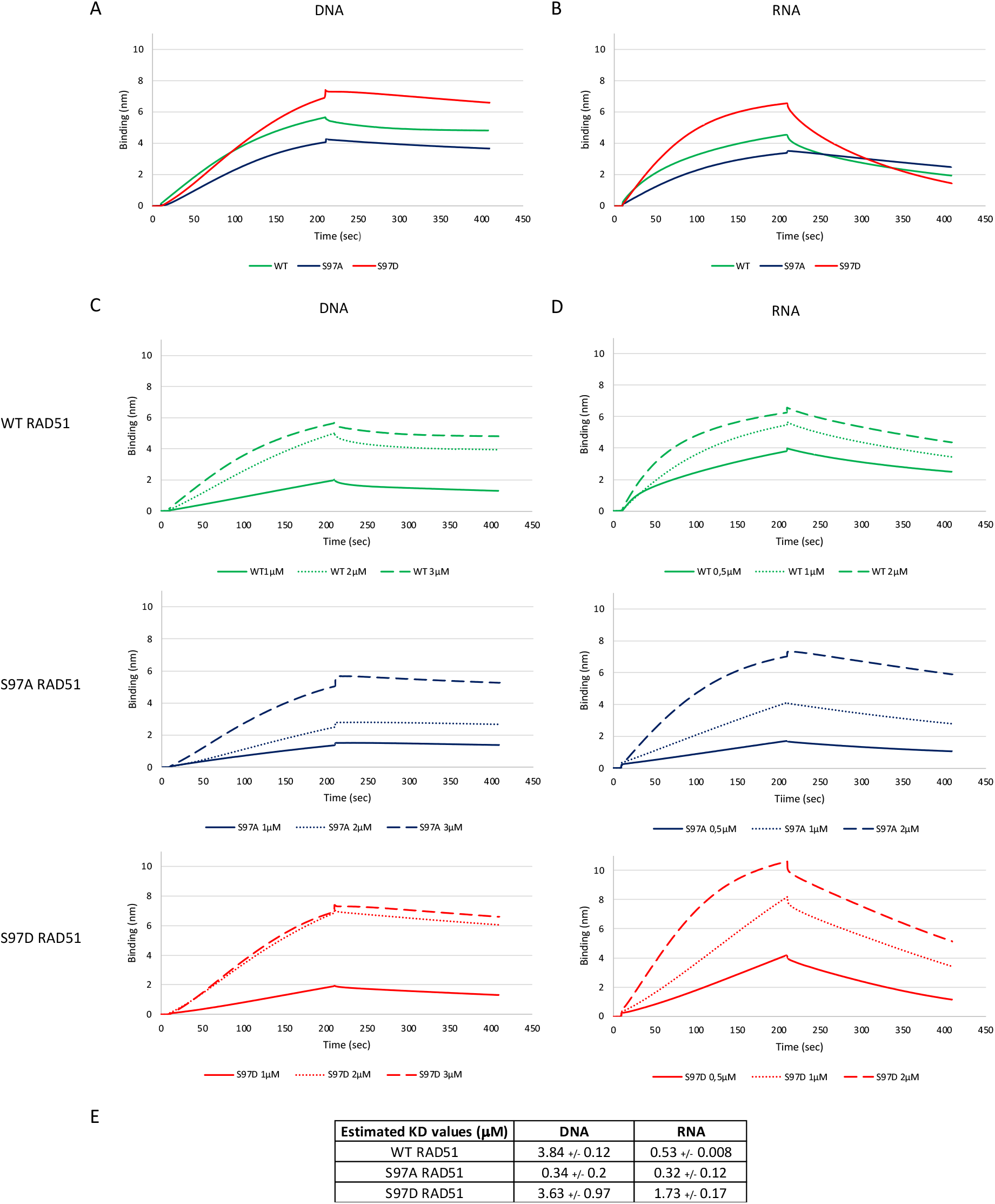
RAD51 phosphorylation mimetic on Ser97 affects its binding to RNA. **A, B**: 3 μM of WT, S97D and S97A RAD51 recombinant proteins were used to evaluate the effect of RAD51 phosphorylation on its binding to 33 nt sized single strand DNA or RNA, in 2 mM ATP final concentration, by using the Blitz technology. **C,D**: Different concentrations, from 0.5 to 3 μM, of WT, S97A and S97D RAD51 recombinant proteins were used to characterize their DNA and RNA binding parameters. **E**: Resulting KD values estimations of WT, S97A and S97D RAD51, binding affinity to DNA and RNA determined from n=4 experiments. Data are means +/− sd.

We show in the figure 7A, the effects of the Ser97 mutations on ssDNA binding capacity. In comparison to the WT RAD51, the S97D RAD51 phosphomimetic mutant showed different association and dissociation curves, with a higher binding capacity and a slightly more rapid dissociation profile. However, the resulting KD evaluation (that takes into account the dissociation/association parameters) showed that the S97D mutant has a ssDNA binding affinity similar to that of the WT (with a KD=3.63 μM for the S97D-RAD51 *vs* a KD=3.84 μM for the WT RAD51). Concerning, the S97A mutant comparison to the WT RAD51, we observed modified association and dissociation curves, with a lower binding capacity and a slower dissociation profile. The resulting KD evaluation showed a higher affinity to ssDNA for the S97A mutant (with a KD=0.34 μM for the S97A-RAD51 *vs* a KD=3.84 μM for the WT RAD51). We can conclude that in our *in vitro* conditions, the Ser97 phosphorylation has no impact on RAD51 affinity for ssDNA, while the S97A mutation provokes a 10-fold higher ssDNA binding affinity. This effect of the S97A mutation will be discussed later.

We next evaluated RAD51’s ability to bind RNA, and the effects of the Ser97 mutations on this ssRNA binding capacity. First, we can see in the figure 7B, that WT and mutants RAD51 are all able to bind RNA. In comparison to the WT RAD51, the S97D mutant showed very different association and dissociation profiles with a higher binding capacity and a faster dissociation profile. These marked differences result in a lower ssRNA binding affinity for the S97D mutant in comparison to the WT RAD51, (with a KD=1.73μM for the S97D *vs* a KD=0.53 μM for the WT RAD51).

Concerning the comparison between the S97A mutant, and the WT RAD51, it shows similar association curves with comparable binding capacities but different dissociation curves. Indeed, the S97A mutant shows a slower dissociation profile. Logically, the resulting KD evaluation showed a slightly higher affinity for ssRNA for the S97 mutant (with a KD=0.32 μM for the S97A *vs* a KD=0.53 μM for the WT RAD51).

All these data showed that RAD51 binds DNA and RNA. The Ser97 mutants highlight the impact of the Ser97 residue as a major modulator of RAD51 affinity for RNA and DNA.

## DISCUSSION

In this work, we identified a new phosphorylation of RAD51 recombinase, the key player in the Homologous Recombination DNA repair pathway. This *in vitro* Aurora A mediated phosphorylation targets the Subunit Rotation Motif of RAD51 at the Ser97 residue. This region is a putative anchoring platform for many other proteins and remains accessible at the surface of the RAD51 nucleofilament. The RecA family SRM, with is evolutionarily conserved, is described as being important in the 3D structure of RecA monomers and is crucial for regulating RecA enzymatic activity ^28^. According to these results, we showed here that the Ser97 phosphorylation has a strong effect on *in vitro* RAD51 D-loop activity, self-association and RNA binding affinity. We showed that RAD51 is an RNA binding protein and that its Ser97 phosphorylation affects strongly its binding to RNA, while it has no effect on its affinity for DNA (WT KD= 3.84 μM and S97D KD=3.63 μM). Indeed, blitz experiments showed that the phosphomimetic RAD51 dissociation from ssRNA is more rapid than that of the WT RAD51 and it lowers its affinity for RNA by a 3-fold factor (KD=1.73μM for the S97D *vs* KD=0.53μM for the WT). This result may seem in contradiction with the *in cellulo* observed presence of PSer97-RAD51 within the nuclear Speckles, but we must consider the difference between the *in vitro* and *in cellulo* conditions where the presence of many partners and PTM of RAD51 can regulate its RNA binding and/or dissociation. Moreover, it is interesting to notice that the S97D phosphomimetic mutant, has a higher binding affinity for ssRNA (KD=1.73 μM) than for ssDNA (KD=3.63μM). Even though the S97D mutant has the same ssDNA affinity than the WT RAD51, it has a higher D-loop activity. The D-loop assay, being a combined evaluation of RAD51/DNA interactions and RAD51 strand exchange activity, we can hypothesize that RAD51 strand exchange activity is enhanced by its phosphorylation on the Ser97 residue. Concerning the S97A mutant comparison to the WT RAD51, we observed that he has higher affinities for both ssDNA (KD= 0.34 μM) and ssRNA (KD= 0.32 μM) with the same affinity for both RNA and DNA. We also observed that the S97A mutation has an effect on RAD51 D-loop activity. This enhanced D-loop activity may be due to a better ssDNA binding or a better strand exchange activity. Even if these results are *in vitro* and do not include physiological regulation of RAD51 activity, they highlighted the importance of the Subunit Rotation Motif and in particular the Ser97 residue in regulating RAD51’s activity and affinity for nucleic acids.

Using a specifically engineered antibody, we also showed that this PSer97-RAD51 does indeed exist *in cellulo*. During RAD51 siRNA experiments, the diminution of RAD51 protein level (63%) is higher than that of the PSer97RAD51 (51%). This could be linked to the PTM effects on proteins half-life. Concerning RAD51, little is known about the effect of its PTMs on its half’s life modulation. Genetic experiments should be performed, using CRISPR/Cas9 technology, in order to answer if the Ser97 phosphorylation affects RAD51 half-life.

Concerning the kinase implicated in this *in cellulo* phosphorylation, we showed that Aurora A overexpression drives a significant enhancement of the Ser97 phosphorylation of RAD51, thus showing that Aurora A is implicated in this new PTM of RAD51. Use of an aurora A inhibitor (alisertib) showed an effect only in the HCC1806 cell line and not in the HeLa cells. These cell line-dependent results points to possible redundancy and illustrate the miss-regulations that occur in different cancers. Therefore, it is possible that other kinases could also phosphorylate RAD51 Ser97 residue. Concerning the characterization of this phosphorylation, we showed that it is decreased after DNA damage induction. It is interesting to notice here that Aurora A kinase activity has been described as decreasing after DNA damage induction ^3^. On the contrary, Aurora A overexpression has been described as inhibiting RAD51 recruitment to DNA breaks without affecting γ-H2AXfoci formation. This inhibition of RAD51 recruitment necessitates Aurora A kinase activity and led to a decrease of HR mediated DNA repair ^29^.

Surprisingly, immunofluorescence experiments showed that the Ser97 phosphorylation is distributed in nuclear foci structures that are not colocalized with γ-H2AX foci. Using a commercial antibody against RAD51, we indeed found that there are some RAD51 foci that are not colocalized with γ-H2AX. Our results revealed an atypical sub-cellular location for RAD51, linked to its Ser97 phosphorylation. Using immunofluorescence and confocal microscopy, we identified these PSer97 foci and showed that they are located within the Nuclear Speckles. It is described in the literature that the use of the splicing inhibitor plaB has an effect on NS activity but keep their structure maintained with an enhancement of their size ^30^. After PlaB treatment, we observed that PSer97-RAD51 foci were still within the Nuclear Speckles even if a slight decrease of the colocalization coefficient was observed. Then, RAD51 recruitment to the NS is independent of a functional splicing activity. Using transitory transfection experiments, we showed that exogenous RAD51 maintained in the cytoplasm had less effect on Nuclear Speckles number, than exogenous RAD51 translocated in the nucleus. Indeed, in HeLa cells, in addition to the decrease of NS number per cell we also observed cells without NS. This phenotype was particularly observed for the non-phosphorylable S97A-RAD51 overexpression. Meaning that the presence of exogenous non-phosphorylable RAD51 in the nucleus is deleterious for Nuclear Speckles. While in HCC1806 cell line, exogenous non-phosphorylable RAD51 maintained in the cytoplasm did not produce this extreme phenotype. The role of the Ser97 residue phosphorylation in RAD51/RNA interaction and NS colocalization needs to be pursued. For that purpose, a comparative analysis of PSer97-RAD51 *vs* RAD51 partners must be performed to understand the regulation of RAD51 binding to RNA within the Nuclear Speckles. Moreover, RAD51 IP showed a co-immunoprecipitation of Sc35. Taken together, our results allow us to consider a possible role of RAD51 in these splicing machineries: a new RAD51 function beyond DNA repair needs to be explored.

RAD51 binding to RNA has also been described by the work of Feretzaki et al. in the context of telomeric metabolism in which they showed that RAD51 is essential for the *in cellulo* recruitment of the TERRA RNA on telomeres and the R-loop formation. These results consolidate our finding of a new phosphorylated form of RAD51 that binds RNA and is found in Nuclear Speckles.

While writing this manuscript, we learned of the work by Damodaran that described a link between Aurora A activity and the splicing machinery. They showed that overexpressed Aurora A is partially colocalized with Sc35, within the Nuclear Speckles, and that Aurora A inhibition provokes splicing alterations. They also showed that Aurora A interacts with and phosphorylates splicing factors ^31^. These data strongly reinforce ours and particularly the implication of Aurora A in the *in cellulo* RAD51 Ser97 phosphorylation.

The DDR requires the orchestrated recruitment of different proteins intervening in chromatin remodeling, cell signaling, and DNA repair. This work provides a basis for further studies exploring more precisely the possible role of the RAD51 DNA repair factor within the Nuclear Speckles’s structure or function, and reveals a new link between DNA repair and alternative splicing regulation. Understanding the regulation of pre mRNA splicing during the DDR is of particular interest as we know that splicing regulation in the context of chemo/radio-therapies is implicated in the development of treatment resistances ^32 26^

## AUTHORS CONTRIBUTION

M.A. made the Cpt, Ali and PlaB cell treatments and WB experiments, Cell-cycle analysis, IP, siRNA experiments.

P.K.Y. made polymerization assays, Blitz DNA and RNA assays. NA made Blitz assay analysis.

V.P.M and G. C made cell culture and protein extractions. A.D. made D-loop assay. D.M made recombinant protein production. HBM conceived, supervised all the study and wrote the manuscript. F.F as the team head made that possible.

## AKNOWLEDGMENTS

This work was financed by the SATT grand-Ouest valorisation.

