## Supplemental figures for "Aurora A mediated new phosphorylation of RAD51 is observed in Nuclear Speckles"

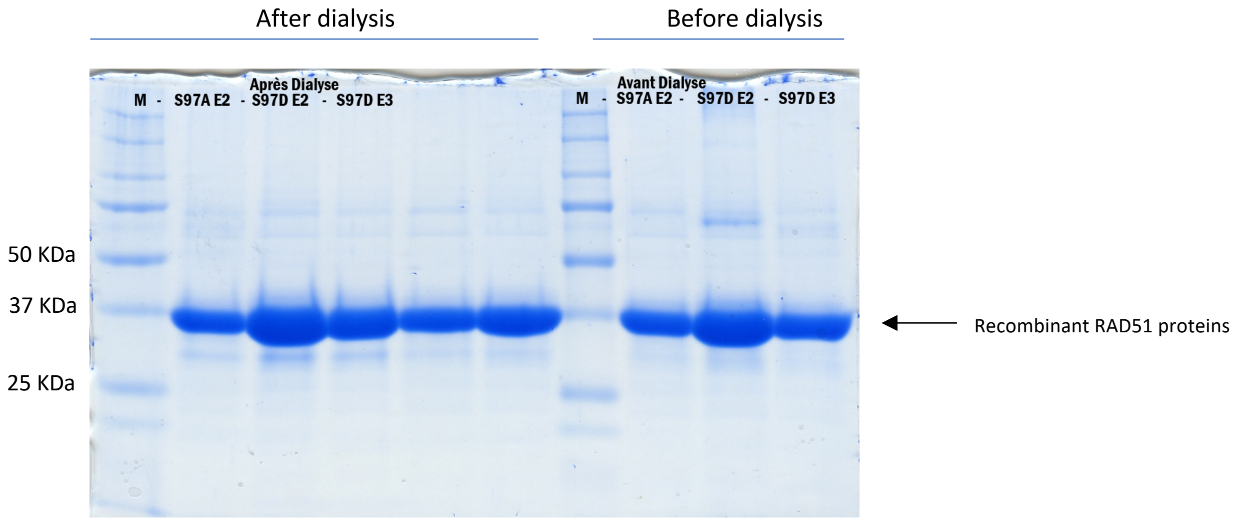


**Supplemental Figure 1: Coomassie Gels of WT, S97A and S97D RAD51 recombinant proteins**
Coomassie blue stained SDS-PAGE showing the recombinant mutant RAD51 proteins.
S97A E2 means S97A-RAD51 Elution N°2. S97D E2 means S97D-RAD51 Elution N°2. S97D E3 means S97D-RAD51 Elution N°3.

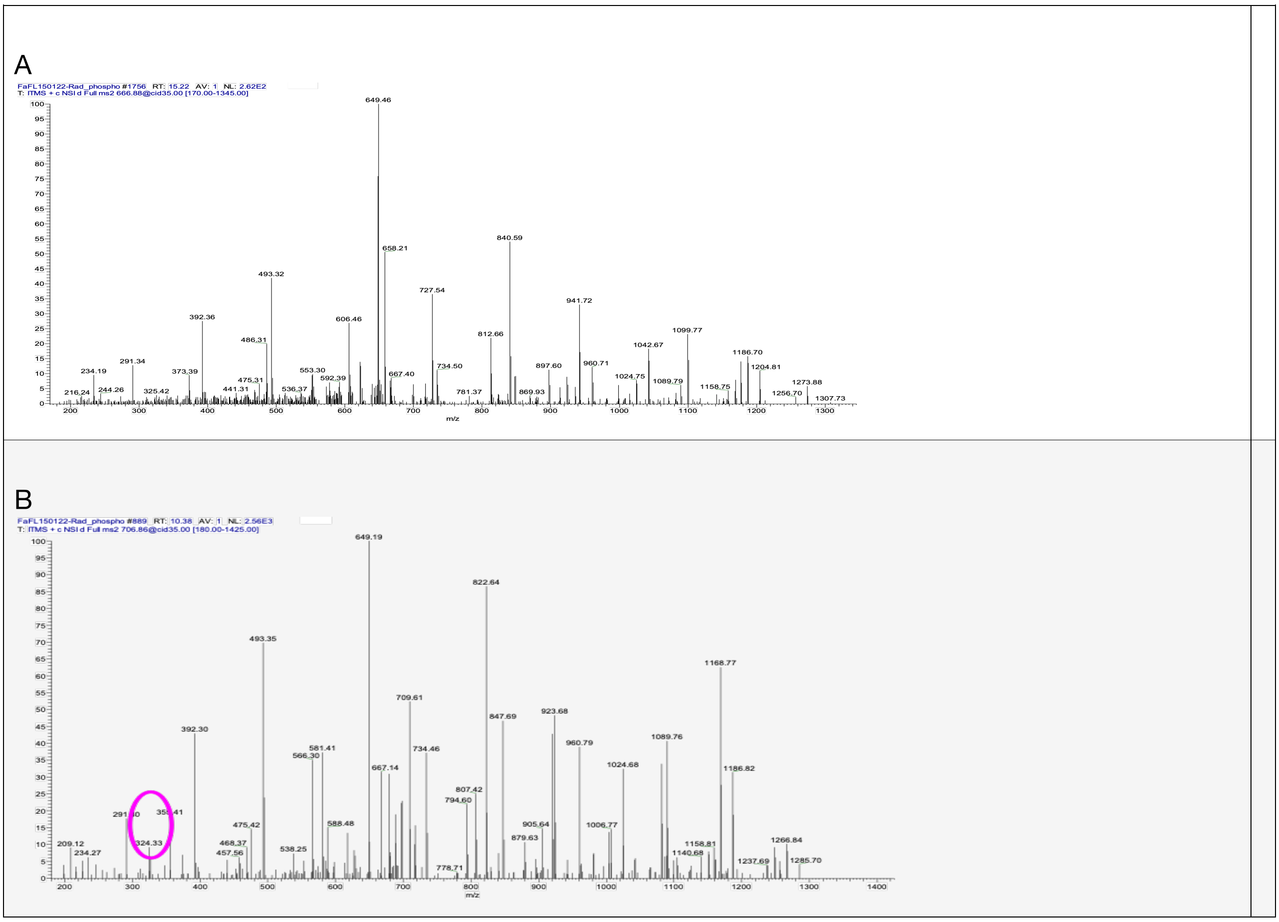


**Supplemental figure 2: Mass spectra of the control and Aurora A-phosphorylated RAD51**
A: mass spectra of the non phosphorylated RAD51 recombinant protein. B: mass spectra of the Aurora A phosphorylated RAD51 recombinant protein. The purple circle identified the phosphorylated residue.

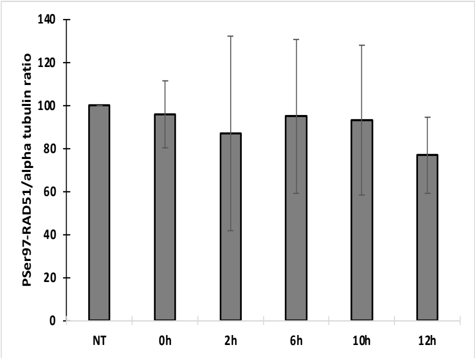


**Supplemental Figure 3: PSer97RAD51/tubulin ratio throughout cell cycle progression**
The PSer97/tubulin ratio was calculated during the cell cycle progression, from n=3 experiments.

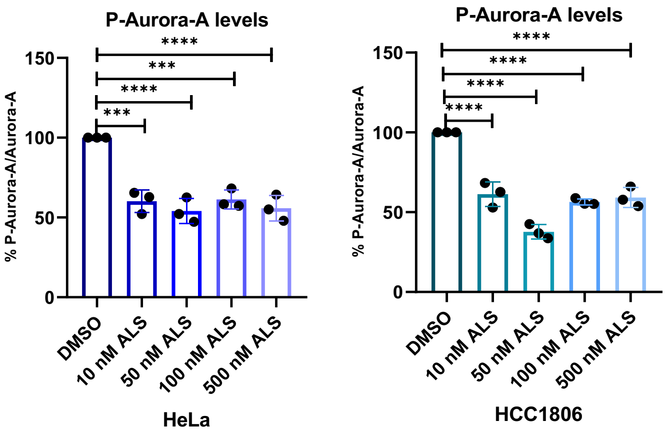


**Supplemental Figure 4: Alisertib efficacy on phosphoThr288Aurora A**

Different Alisertib concentrations were used in HeLa and HCC1806 cell lines, for 24 h treatment. Whole cell extracts were used to perform WB and evaluate the effect on the pThr288, auto-phosphorylated form of Aurora A kinase. Statistical analysis using paired t test were performed, n=3. p value ($\alpha$= 0,05: *, $\alpha$= 0,001: **, $\alpha$= 0,001: ***).
